# Normalisr: normalization and association testing for single-cell CRISPR screen and co-expression

**DOI:** 10.1101/2021.04.12.439500

**Authors:** Lingfei Wang

## Abstract

Single-cell RNA sequencing (scRNA-seq) provides unprecedented technical and statistical potential to study gene regulation but is subject to technical variations and sparsity. Here we present Normalisr, a linear-model-based normalization and statistical hypothesis testing framework that unifies single-cell differential expression, co-expression, and CRISPR scRNA-seq screen analyses. By systematically detecting and removing nonlinear confounding from library size, Normalisr achieves high sensitivity, specificity, speed, and generalizability across multiple scRNA-seq protocols and experimental conditions with unbiased P-value estimation. We use Normalisr to reconstruct robust gene regulatory networks from trans-effects of gRNAs in large-scale CRISPRi scRNA-seq screens and gene-level co-expression networks from conventional scRNA-seq.

## Background

Understanding gene regulatory networks and their phenotypic outcomes forms a major part of biological studies. RNA-sequencing (RNA-seq) has received particular popularity for systematically screening transcriptional gene regulation and co-regulation. ScRNA-seq provides a unique glance into cellular transcriptomic variations beyond the capabilities of bulk technologies, enabling analyses such as single-cell differential expression (DE) [1], co-expression [2, 3], and causal network inference [4, 5] on cell subsets at will. Regarding and manipulating each cell independently, especially in combination with CRISPR technology [6, 7], scRNA-seq can overcome the major limitations in sample and perturbation richness of bulk studies, at a fraction of the cost.

However, cell-to-cell technical variations and low read counts in scRNA-seq remain challenging in computational and statistical perspectives. A comprehensive benchmarking found single-cell-specific attempts in DE could not outperform existing bulk methods such as edgeR [8]. Generalized linear models (*e.g*. [9, 1]) are difficult to generalize to single-cell gene co-expression, leaving it susceptible to expression-dependent normalization biases [10]. CRISPR scRNA-seq screen analysis can be regarded as multi-variate DE, but existing methods (e.g. scMageck [11] and SCEPTRE [12]) take weeks or more on a modern dataset and do not account for off-target effects.

A potential unified framework for single-cell DE, co-expression, and CRISPR analysis is a two-step normalization-association inferential process (Fig. 1a). The normalization step (e.g. sctransform [13], bayNorm [14], and Sanity [15]) removes confounding technical noises from raw read counts to recover the biological variations. The subsequent linear association step has been widely applied in the statistical analyses of bulk RNA sequencing and micro-array co-expression [16], genome-wide association studies (GWAS) [17, 18], expression quantitative trait loci (eQTL) [19], and causal network inference [20, 21]. Linear association testing presents several advantages: (i) exact P-value estimation without permutation, (ii) native removal of covariates (*e.g*. batches, house-keeping programs, and untested gRNAs) as fixed effects, and (iii) computational efficiency. However, existing normalization methods were designed primarily for clustering. Sensitivity, specificity, and effect size estimation in association testing are left mostly uncharted.

**Figure 1:**
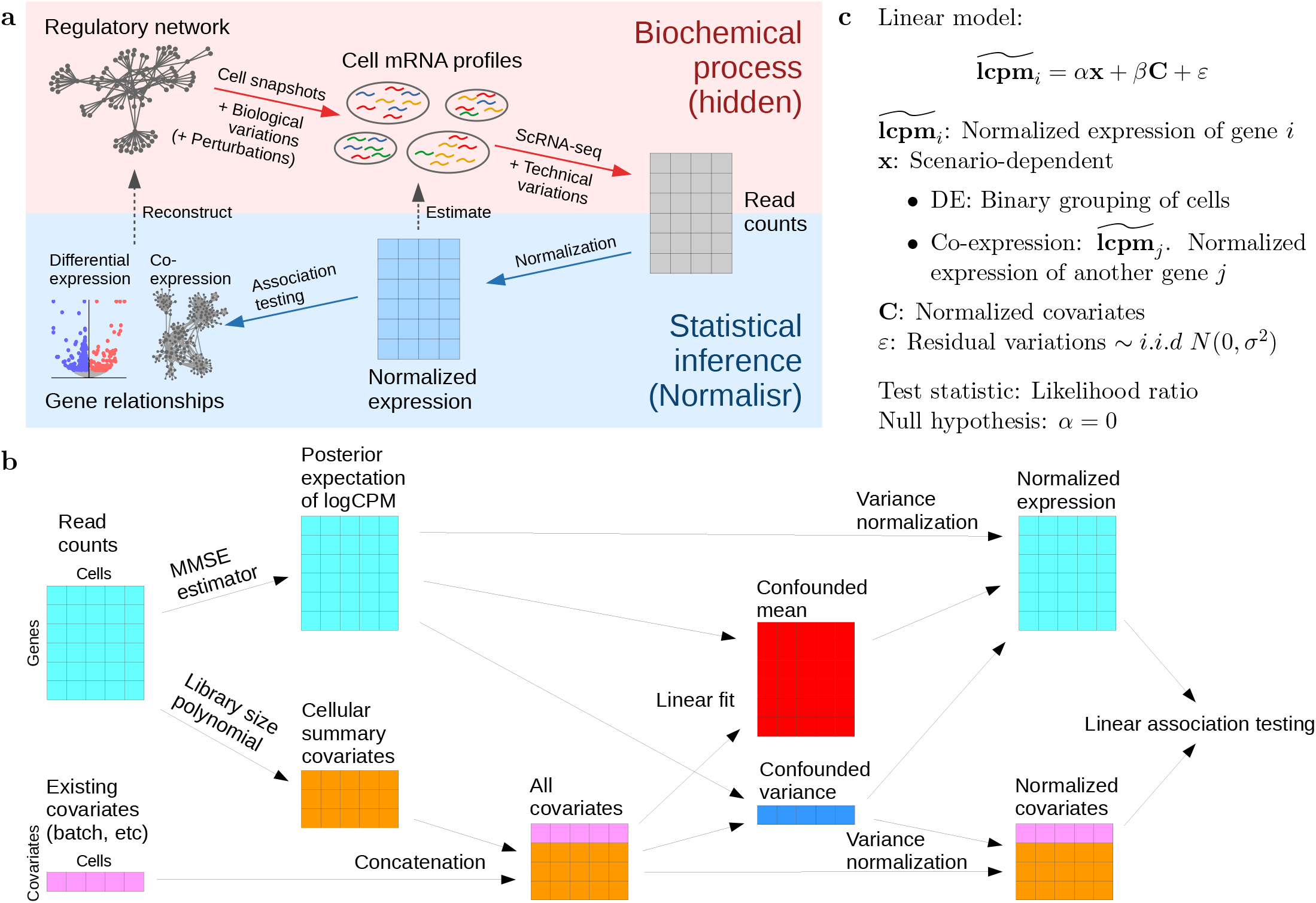
Normalisr overview. **a** Schematics of biochemical and inferential processes (top to bottom). Normalisr aims to remove technical variations and normalize raw read counts to estimate the pre-measurement mRNA abundances. Then they can be directly handled by conventional statistical methods, such as linear models, to infer gene regulation and to unify different analyses. **b** Normalization step (in **a**) of Normalisr (left to right). Normalisr starts by computing the expectation of posterior distribution of mRNA log proportion in each cell with MMSE. Meanwhile, Normalisr appends existing covariates with nonlinear cellular summary covariates. Normalisr then normalizes expression variance by linearly removing the confounding effects of covariates on log variance. The normalized expression and covariates are ready for downstream association testing with linear models, such as differential and co-expression (in **c**). **c** Association testing step (in **a**) of Normalisr, which uses linear models to test gene differential and co-expression.

Here, we present Normalisr, a normalization-association framework for statistical inference of gene regulation and co-expression in scRNA-seq (Fig. 1a). The normalization step estimates the pre-measurement mRNA frequencies from the scRNA-seq read counts (*e.g*. unique molecular identifiers or UMIs) and regresses out the nonlinear effect of library size on expression variance (Fig. 1b). The association step utilizes linear models to unify case-control DE, CRISPR scRNA-seq screen analysis as multivariate DE, and gene co-expression network inference (Fig. 1c). We demonstrate Normalisr’s applications in two scenarios: gene regulation screening from pooled CRISPRi CROP-seq screens and the reconstruction of transcriptome-wide co-expression networks from conventional scRNA-seq.

## Results

### Normalisr overview

Normalisr starts normalization with the minimum mean square error (MMSE) estimator of log counts-per-million (Bayesian logCPM). Briefly, this Bayes estimator computes the posterior expectation of log mRNA abundance for each gene in each cell based on the binomial (approximate of multinomial) mRNA sampling process [10, 22] (Methods). This naturally prevents the artificial introduction and choice of constant in log(CPM+constant). We also intentionally avoided imputations that rely on information from other genes that may introduce spurious gene inter-dependencies, or from other cells that assume whole-population homogeneity and consequently risk reducing sensitivity (see below).

To account for potential technical confounders, we introduced two previously characterized cellular co-variates: the number of zero-read genes [1] and log total read count [13]. These two covariates comprise the complete set of unbiased cellular summary statistics up to first order (as *L*^0^ and *L*^1^ norms respectively) without artificial selection of gene subsets. Restricting to unbiased covariates minimizes potential interference with true co-expressions. The number of zero-read genes also provides extra information in the read count distribution for each cell and allows for a regression-based automatic correction for distribution distortions. This approach aims for the same goal as quantile normalization but is not restricted by the low read counts in scRNA-seq.

We modeled nonlinear confounding effects on gene expression with Taylor polynomial expansion. We iteratively introduced higher order covariates to minimize false co-expression on a synthetic co-expression-free dataset, using forward stepwise feature selection that is restricted to series expansion terms whose all lower order terms had already been included. The set of nonlinear covariates were pre-determined as such and then universally introduced for all datasets and statistical tests. We first linearly regressed out these covariates’ confounding effects at the log variance level [23]. Their mean confounding were accounted for in the final hypothesis testing of linear association (Fig. 1bc). Finally, P-values were computed from Beta distribution of *R*^2^ in exact conditional Pearson correlation test (equivalent of likelihood ratio test) without permutation.

### Normalisr detected nonlinear technical confounders with restricted forward stepwise selection

We downloaded a high multiplicity of infection (high-MOI) CRISPRi Perturb-seq dataset of K562 cells [6] based on 10X Genomics to determine the nonlinear confounding of cellular summary covariates. From this, we generated a synthetic null scRNA-seq dataset to mimic the gRNA-free K562 cells but without any co-expression, with log-normal and multinomial distributions respectively for biological and technical variations (Fig. S1, Methods). (For exact gene, cell, and gRNA counts here and onwards, see Table S1.)

Using the cellular summary covariates as the basis features and optimizing towards a uniform distribution of (conditional) Pearson co-expression P-values, we confirmed that both the log total read count and the number of zero-read genes in a cell strongly confound single-cell mRNA read count (Fig. S2). Moreover, we identified strong nonlinear confounding from the square of log total read count. Together, these three extra covariates were suffice to recover the uniform null distribution of co-expression P-values. These predetermined nonlinear cellular covariates allowed Normalisr to perform hypothesis testing without any parameter or permutation in subsequent evaluations and applications.

### Evaluations in single-cell DE and co-expression

We performed a wide range of evaluations of Normalisr and other methods with the Perturb-seq dataset. We partitioned cells that did not detect any gRNA into two random groups 100 times to evaluate the null P-value distribution of single-cell DE. To detect expression-dependent null P-value biases, we grouped genes into 10 equally sized subsets from low to high expression and compared their null P-values against the uniform distribution with the Kolmogorov-Smirnov (KS) test (Methods). Normalisr recovered uniform distributions of null DE P-values at all expression levels, as did Sanity, bayNorm (with ‘mode’ or ‘mean’ options), and sctransform that were followed by the same linear model (Fig. 2ab, Fig. S3). In contrast, edgeR and MAST, the best single-cell DE methods benchmarked in [8], had expression-dependent, 0- or 1-biased null P-values, which would lead to high false positive and false negative rates in application. Imputation methods such as MAGIC [24] and DCA [25] could not recover a uniform distribution either. Normalisr was additionally much faster than most other methods tested, and over 2,000 times than edgeR and MAST (Fig. 2c).

**Figure 2:**
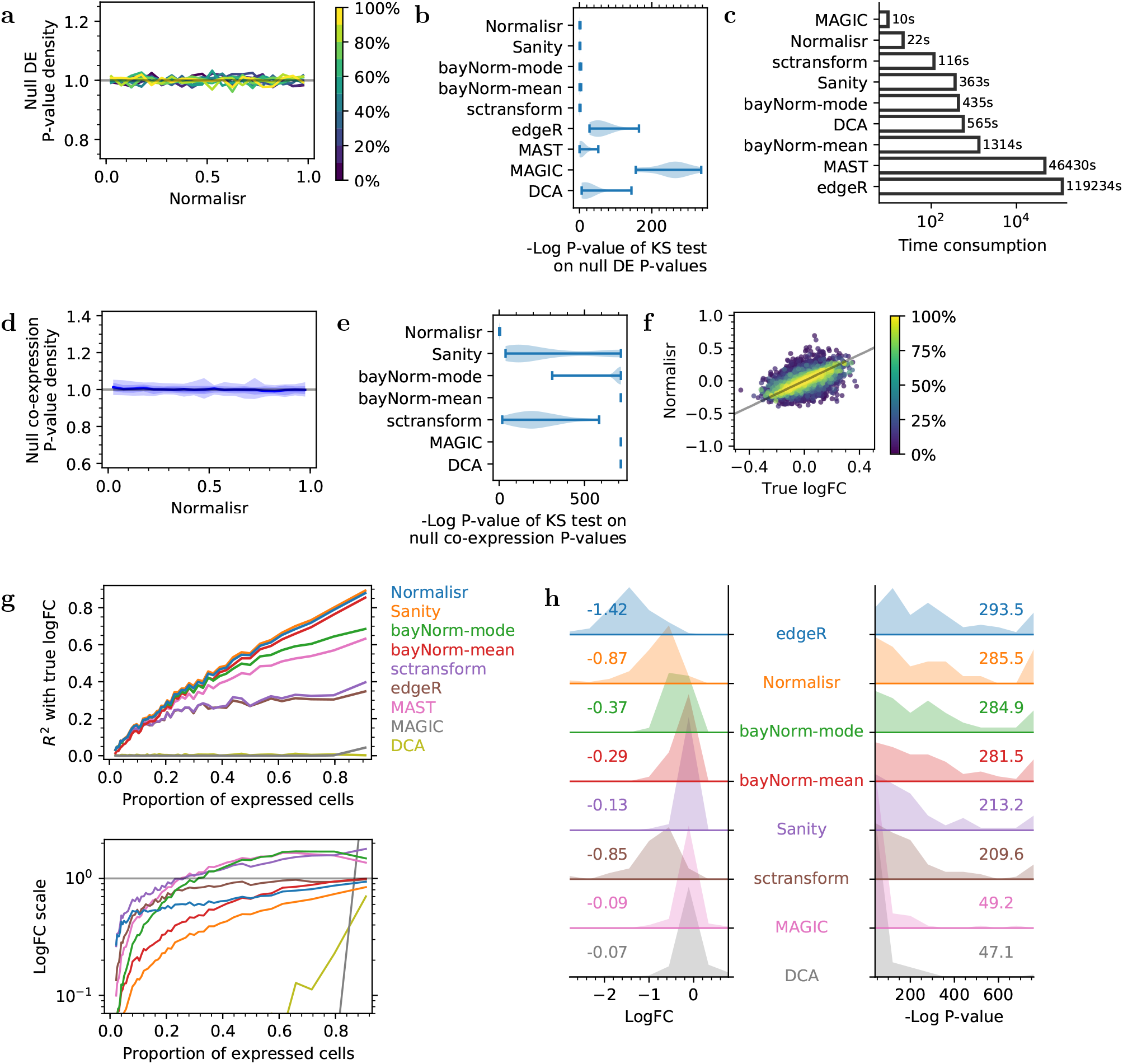
Normalisr achieved high sensitivity, specificity, and speed in single-cell DE and co-expression. **a** Normalisr had uniformly distributed null P-values in single-cell DE (X) as shown by histogram density (Y). Genes were evaluated in 10 equally sized and separately colored bins stratified by expression (proportion of expressed cells). Gray line indicates the expected uniform distribution for null P-value. **b** Normalisr, Sanity, bayNorm, and sctransform recovered uniformly distributed null P-values for DE, as measured by the violin plot of P-values of KS test (X) on null P-values separately for each gene bin (in **a**). Bars show extrema. **c** Normalisr was much faster than other normalization methods. **d** Normalisr had uniformly distributed null P-value in single-cell co-expression (X) from synthetic data, as shown by histogram density (Y). Genes were split into 10 equal bins from low to high expression. The null P-value distribution of co-expression between each bin pair formed a separate histogram curve. Central curve shows the median of all histogram curves. Shades show 50%, 80%, and 100% of all histograms. Gray line indicates uniform distribution. **e** No other method tested could recover uniformly distributed null P-values for co-expression, as measured by the violin plot of P-values of KS test (X) on null P-values separately for each gene bin pair (in **d**). Bars show extrema. **f** Normalisr accurately recovered logFCs (Y) when compared against the synthetic ground-truth (X) for genes from low to high expression (color). Gray line indicates X=Y. **g** Normalisr accurately recovered logFCs with low variance (top, Y as *R*^2^) and low bias (bottom, Y as regression coefficient; horizontal gray line indicates bias-free performance) when evaluated against synthetic ground-truth with a linear regression model separately for genes grouped from low to high expression (X) on logFC scatter plots (**f** and Fig. S5). **h** Normalisr was the most sensitive (right, X) and recovered the most accurate logFCs (left, X) among normalization and imputation methods in the CRISPRi Perturb-seq experiment. Histogram shades show distributions of inferred DE statistics of the targeted gene between KD and unperturbed cells (positive controls with varying strengths). Numbers indicate mean values. Stronger P-values and logFCs closer to edgeR indicates higher sensitivity and accuracy.

To detect potential incomplete removal of technical confounding, we compared the distribution of co-expression P-values from the synthetic null dataset. Normalisr recovered uniformly distributed P-values irrespective of expression and better than log(CPM+1) with or without covariates (Fig. 2de, Fig. S4), suggesting the necessities of both the Bayesian logCPM and the nonlinear cellular summary covariates. Sanity, bayNorm, and sctransform could not fully account for technical confounding and incurred distorted null P-value distributions. Imputation methods inflated gene-gene associations because of their inherent reliance on gene relationships. Normalisr accounted for technical confounding and correctly recovered the absence of co-expression.

The synthetic dataset recorded the biological ground-truths of cell mRNA profiles (Fig. 1a) and allowed for evaluations in log Fold-Change (logFC) estimation. For this, we decomposed logFC estimation errors of each method into bias and variance with linear regression against the ground-truth logFC. Bias indicates the overall under- or over-estimation errors in logFC scale as the deviation of regression coefficient from unity. Variance represents uncertainties in the estimation and was quantified with *R*^2^ of the regression. Normalisr had the lowest variance among normalization and imputation methods and accurately estimated logFCs across all expression levels (Fig. 2fg, Fig. S5). BayNorm underestimated the logFCs of lowly expressed genes, and Sanity underestimated the logFCs regardless of expression. Their prior distributions aggregate information across cells under the implicit assumption of population homogeneity, which may over-homogenize gene expression and consequently underestimate logFCs, especially for lowly expressed genes whose fewer reads possessed less information to overcome the prior. On the other hand, edgeR recovered the most accurate logFCs for moderately and highly expressed genes but was susceptible to large bias for lowly expressed genes and large overall variance. Normalisr accurately estimated logFCs with low bias and variance for genes across all expression levels.

We then assessed the sensitivity of different methods on single-cell DE. In the absence of a high-quality gold standard dataset, we focused on the intended CRISPRi effects on targeted gene expression by comparing cells infected with each single gRNA against uninfected cells (excluding cells infected with any other gRNA). Normalisr achieved comparable sensitivity in terms of P-value with edgeR but was more sensitive on highly expressed genes and less so on lowly expressed genes (Fig. 2h, Fig. S6), possibly due to edgeR’s expression-dependent biases on the null dataset (Fig. S3). Normalisr also estimated logFCs more accurately than other normalization or imputation methods when using edgeR as a proxy for ground-truth, which performed best in the synthetic dataset. Sanity and sctransform suffered sensitivity losses while Sanity and bayNorm underestimated logFC, similar to observations on the synthetic dataset. Imputation methods lost the major effects of CRISPRi. Normalisr recovered the strongest P-values and the most accurate logFCs among normalization and imputation methods.

Our evaluation results were reproducible on a MARS-seq dataset [26] of dysfunctional T cells from frozen human melanoma tissue samples (Fig. S7) that is further explored for co-expression later. Particularly, both the nonlinear covariate set and Normalisr’s performance were generalizable across distinct sequencing methodologies, sequencing depths, and experimental conditions. Normalisr provides a normalization framework with improved sensitivity, specificity, and efficiency to allow linear association testing for single-cell differential and co-expression that are consistent across multiple scRNA-seq protocols and experimental conditions.

### False positives from gRNA cross-associations in high-MOI CRISPR screens

High-MOI CRISPRi systems are highly efficient screens for gene regulation, gene function [6, 27], and regulatory elements [7]. However, numerous factors may confound gRNA-mRNA associations and lead to high false positive rates (FPR), such as competition between gRNAs for dCas9 or for limited read counts. These competitions may lead to gRNA cross-associations, which incur false positive up- and down-regulations of genes regulated by other gRNAs or their target genes (Fig. 3a).

**Figure 3:**
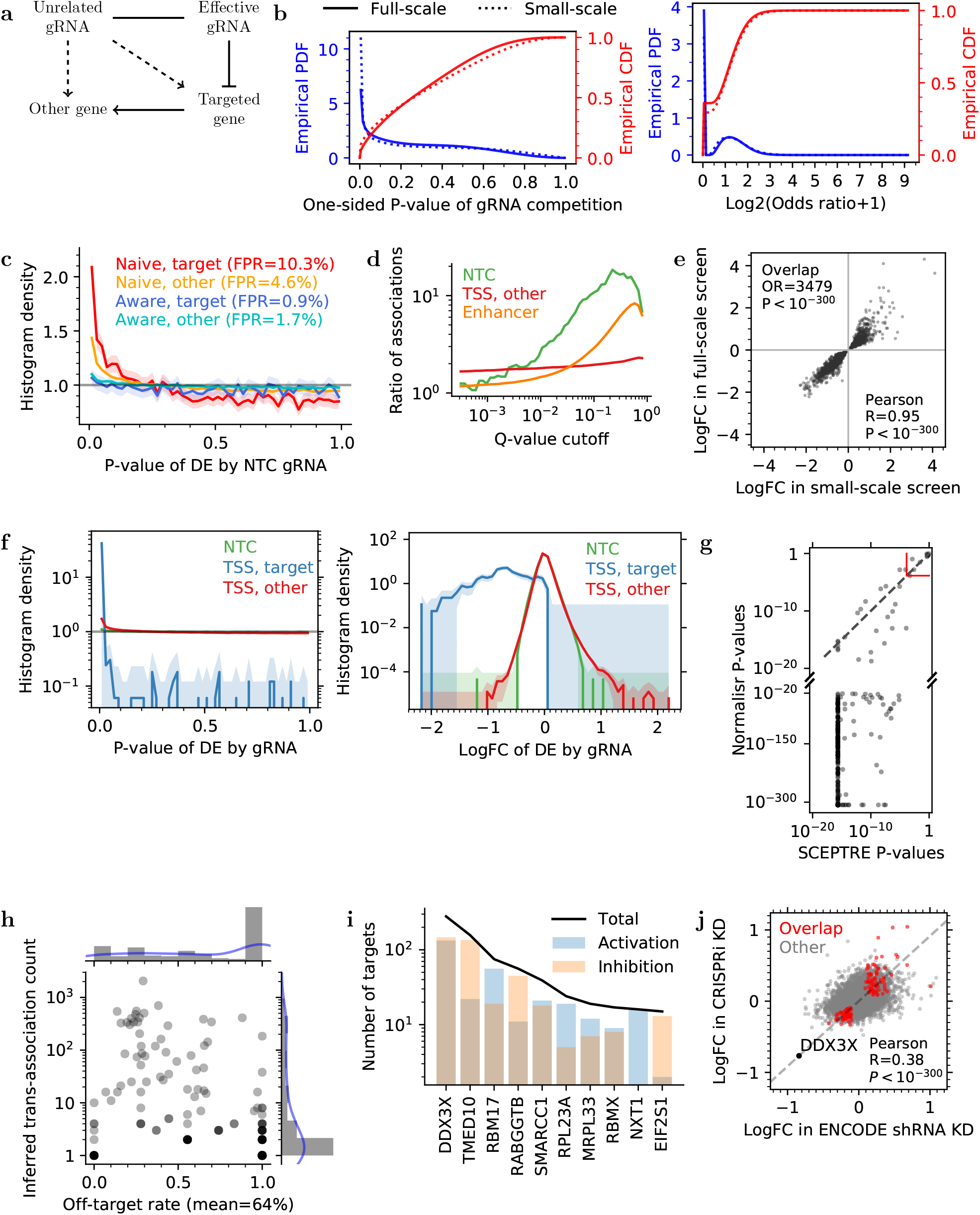
Normalisr detected robust and specific gene regulation from high-MOI CRISPRi systems. **a** Example scenario of false positives of gRNA-gene associations (dashed) arising from negative gRNA cross-associations and true regulation (solid) for genes targeted directly by the effective gRNA or indirectly through the targeted genes (other gene). **b** Detection of different gRNAs were anti-correlated across cells in terms of one-sided hypergeometric P-values (left, X) and odds ratios (right, X) between gRNA pairs. Empirical PDFs and CDFs are shown in blue (left Y) and red (right Y) respectively. **c** Ignoring untested gRNAs increased FPR, as shown by density histogram (Y) of CRISPRi DE P-values from NTC gRNAs (X). P-value histograms and FPRs were computed separately with competition-**naive** (ignoring untested gRNAs) and -**aware** (regarding untested gRNAs as covariates) methods and separately for genes **target**ed by positive control gRNAs at the TSS and **other** genes. **d** Competition-naive DE inflated the number of significant gRNA-gene associations relative to the competition-aware method (Y) at different nominal Q-values (X). **e** Guide RNA-gene associations were highly reproducible among 1,857 significant regulations (dot) between the logFCs in small- (X) and full-scale (Y) CRISPRi screens. **f** Normalisr uncovered gene regulations as secondary effects of TSS-targeting gRNAs, which was significantly stronger than NTCs in terms of histograms (Y) of DE P-value (left, X) and logFC (right, X). **g** Normalisr is more sensitive than SCEPTRE on positive control CRISPRi repressions (dots) in terms of P-values. Dashed gray line indicates equal sensitivity. Solid red line is the significance cutoff by Normalisr or SCEPTRE (Bonferroni *P* < 0.05). **h** Over half of significant trans-associations were potential off-target effects at *Q* < 0.05. The putative off-target rate (X) among the number of inferred trans-associations (Y) is shown for each gene (dot) that is directly targeted by two gRNAs at the TSS. **i** Numbers of inferred targets (Y) of top regulators (X) showed different activation and inhibition preferences. **j** Gene DE logFC by DDX3X knock-down in CRISPRi screen (Y) were confirmed with that from ENCODE shRNA knock-down (X). Overlapped targets are highlighted in red (OR=1.16, hypergeometric *P* = 0.01). Dashed line passes through DDX3X and the origin. **Histograms** Gray lines indicate the expected uniform distribution for null P-value. Shades indicate absolute errors estimated as 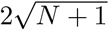, where ***N*** is the count in each bin.

To understand the frequency of gRNA cross-associations, we utilized the enhancer-screening CROP-seq dataset [7] of 207,324 K562 cells post quality control, containing 13,186 gRNAs and 10,877 genes. Cells already containing gRNAs were indeed less likely to include another gRNA (Fig. 3b, Methods), significantly deviating from a random gRNA assignment. This was reproducible for a small-scale screen in the same study.

We then used non-targeting-control gRNAs (NTCs) as negative controls to estimate the elevated FPR from gRNA-gene associations. A competition-naive DE analysis using Normalisr that disregarded other, untested gRNAs lead to a 4.8% overall FPR among NTCs (Fig. 3c, Fig. S8, Storey’s method [28]). Notably, the FPR was significantly higher among genes whose transcription start sites (TSS) were targeted by another gRNA. This supported our hypothesis that false positives were mediated by TSS-targeting gRNAs, because indirect effects through the targeted genes are weaker and harder to detect (Fig. 3a).

### Application to High-MOI CROP-seq screens

Linear models can account for other, untested gRNAs as additional covariates in a competition-aware model. Inspired by [29], we also reduced the covariates, when more than 10,000, to their top 500 principal components as an efficient heuristic solution, which was evaluated to be robust on the small-scale screen (Fig. S9, Methods). This allowed Normalisr to statistically test all cis- and trans-regulations between every gRNA-gene pair in the full-scale screen within a day on a 64-core computer. In comparison, scMAGECK failed to finish within two weeks and SCEPTRE, edgeR, and MAST are projected to need over a year. Normalisr was uniquely efficient for modern-scale CRISPR scRNA-seq screens.

Normalisr successfully recovered NTCs’ improved, near-uniform P-value distribution independent of whether the gene was targeted at the TSS (Fig. 3c, Fig. S8, overall FPR=1.5%). On contrary, the competition-naive method under-estimated the false discovery rate (FDR), reporting over 15 times the gRNA-gene associations for NTC gRNAs and over 4 times for candidate enhancer-targeting gRNAs at nominal *Q* ≤ 0.2 than the competition-aware method (Fig. 3d).

To validate the reproducibility of significant gRNA-gene associations (Bonferroni *P* ≤ 0.05 in each screen), we performed the same inference on the small-scale screen. We found major overlap between the two screens, with highly correlated logFCs and all effect directions matched (Fig. 3e). We also detected unexpected associations of NTC gRNAs with gene expression that were consistent between screens (*P* < 3 × 10^-4^ in each screen and total Bonferroni *P* < 0.1, Table S2). The affected genes were uniformly repressed suggesting potential off-targeting, with the exception of ZFC3H1 that plays a major role in RNA degradation [30]. Normalisr’s regulation inference was reproducible across the screens.

Guide RNAs targeting TSS were regarded as positive controls in [7], and indeed we found 95% (703/738 at *Q* ≤ 0.05) to significantly inhibit the expression of targeted genes with varying efficiencies (Fig. 3f). In comparison with SCEPTRE that was regarded most sensitive[12], Normalisr was even more sensitive at gene level (Fisher’s method) and could identify most (348/357) of these positive controls at a stronger P-value (Fig. 3g). This lead remained apparent even after considering SCEPTRE’s P-value lower bound from resampling.

Despite the dataset’s original design as an enhancer screen, we repurposed it to infer gene regulatory networks by searching for TSS-targeting gRNA→targeted gene→trans-gene relationships via trans-association. With two gRNAs targeting the same TSS, we quantified the proportion of off-target effects by looking for trans-associations that were significant for the weaker gRNA (in association with the targeted gene) but not for the stronger gRNA. We found over half of the inferred trans-associations were likely off-target effects on average (Fig. 3h). The mean off-target rate was significantly reduced at a more stringent threshold (Fig. S10), suggesting that the putative off-target effects were on average weaker than genuine trans-associations.

To account for off-target effects and mediation through nearby genes, we excluded gRNAs that inhibited another gene within 1Mbp from the TSS and gene regulations irreproducible across gRNAs targeting the same TSS or across screens. In total, we recovered 833 high-confidence putative gene regulations (all *Q* ≤ 0.2) that formed a gene regulatory network (Table S3).

Some of the top identified regulators exhibited strong preferences in up- or down-regulation of other mRNAs (Fig. 3i, Table S4). TMED10, *a.k.a* p24*δ*_1_, is responsible for selective protein trafficking at the endoplasmic recticulum (ER)-Golgi interface [31], and indeed its inhibition up-regulated ER- and Golgi-localized genes. Inhibition of RBM17, part of the spliceosome and essential for K562 cells (ranked at 3.8% in CRISPR knock-out screen [32]), modulated cell respiration gene expression. RABGGTB up-regulated vesicle-localized genes, in agreement with its role as a Rab geranylgeranyl transferase subunit [33].

We cross-validated Normalisr’s DE against bulk RNA-seq of K562 cells with DDX3X knock-down and control shRNA vectors from the ENCODE consortium [34, 35] using edgeR (*Q* ≤ 0.05). Despite its limited sample size (2 each) and differently engineered cell lines, we observed an agreement in logFC and significant overlap of DE genes (Fig. 3j). In conclusion, Normalisr detected specific and robust gene regulations in high-MOI CRISPRi screens that could be cross-validated with existing bulk datasets and domain knowledge.

### Normalisr discovered functional gene modules with single-cell co-expression network in dysfunctional T cells

Organisms are evolved to efficiently modulate various functional pathways, partly through the regulation and co-regulation of gene expression [16]. Manifested at the mRNA level, gene co-expression may provide unique insights into gene and cell functions. However, no existing co-expression detection or network inference method could control for false discovery at the single-cell, single-time-point level [5]. As shown with synthetic null datasets, Normalisr can account for nonlinear confounding from library size and therefore control the FDR in co-expression detection.

We used Normalisr to infer single-cell transcriptome-wide co-expression networks in dysfunctional CD8+ T cells in human melanoma MARS-seq [26]. After removal of low-variance outlier samples (9 of 15,537, Fig. S11), co-expression predominantly arose from cell-to-cell variations in housekeeping functions (Fig. S12, Methods). The top 100 principal genes — those with the most co-expression edges — were enriched with cytosolic and ribosomal genes and reflected translational activity differences across cells (Fig. 4a, Table S5). These housekeeping pathways dominated the correlations in mRNA expression and obscured the cell-type-specific pathways and interpretations of the co-expression network. Meanwhile, spike-ins clustered strongly together and with pseudogenes despite the unbiased analysis, suggesting limitations of spike-ins as a gold standard for null co-expression.

**Figure 4:**
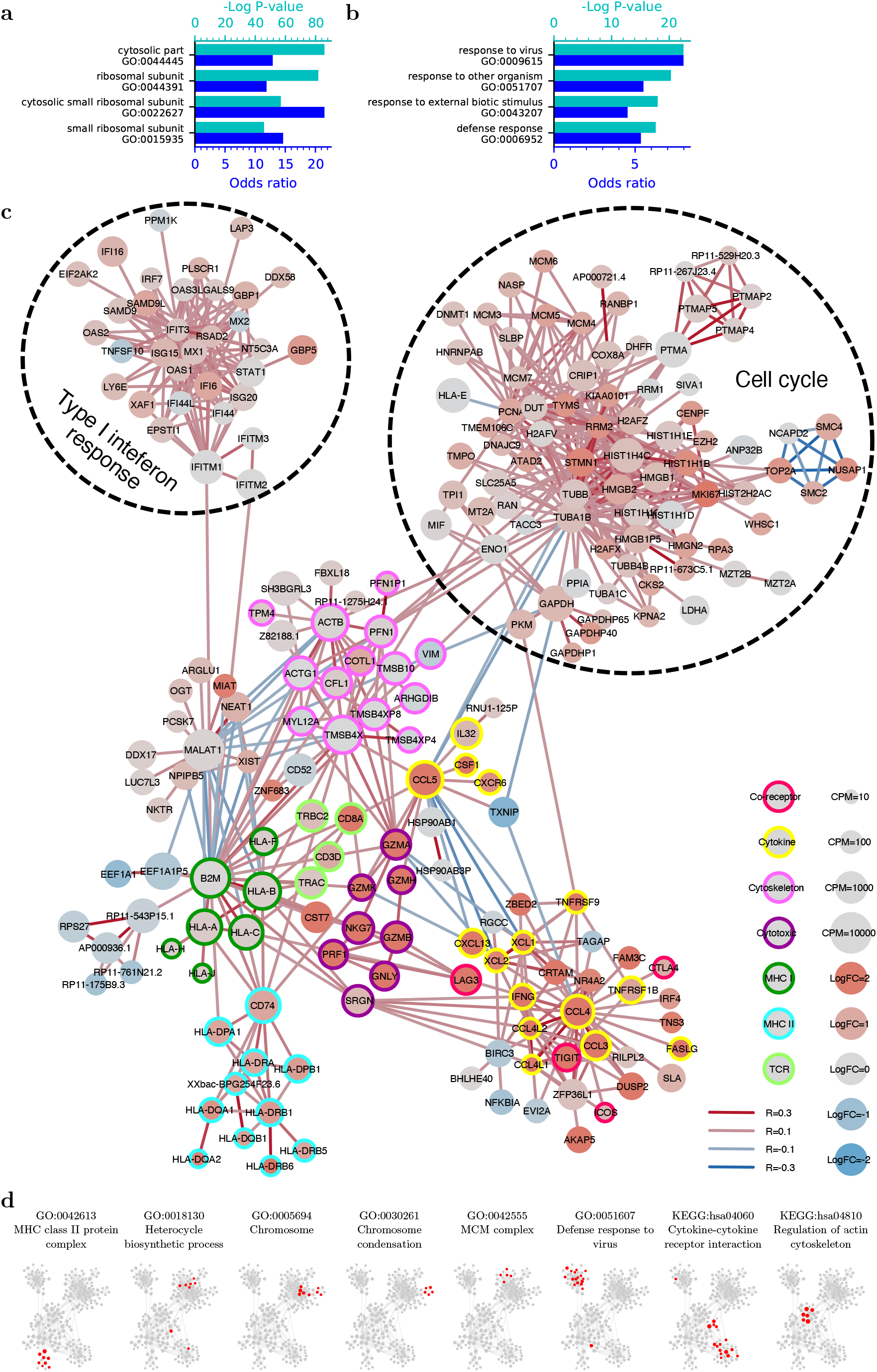
Normalisr revealed gene-level cellular pathways and functional modules in the single-cell co-expression network. **a,b** Single-cell co-expression was dominated by house-keeping programs (**a**), whose iterative removal recovered cell-type-specific programs (**b**), according to the top 4 GO enrichments (Y) of the top 100 principal genes in the co-expression network. P-values and odds ratios are shown in cyan (top X) and blue (bottom X). **c** Single-cell transcriptome-wide co-expression network (major component) after house-keeping program removal highlighted functional gene sets for dysfunctional T cells. Edge color indicates positive (red) or negative (blue) co-expression in Pearson R. Node color indicates DE logFC between dysfunctional and naive T cells. Node size indicates average expression level in dysfunctional T cells. Node border annotates known gene functions. Dashed circles indicate major gene co-expression clusters. **d** Single-cell co-expression recovered cell-type-specific and generic gene functional similarities, according to significant over-abundances of edges between genes in the same GO or KEGG pathway, as highlighted in red.

To focus on cell-type-specific pathways, we developed a sub-routine to remove the high-confidence but generic co-expressions as additional covariates. For this, we identified the strongest GO enrichment from the top 100 principal genes. We then introduced the top principal component (PC) of expression of the genes in this GO category as an additional covariate before re-computing the co-expression statistics and the GO enrichment of top principal genes. Iteratively, we removed the top PCs of GO processes corresponding to ‘cytosolic part’ (reflecting translation) and ‘chromosome condensation’ (reflecting cell cycle). The top GO enrichments subsequently reflected immune system processes, signifying recovery of cell-type-specific co-expressions and also the end of iteration (Fig. 4b, Fig. S12, Table S5).

Meanwhile, we performed single-cell DE between dysfunctional and naive T cells and recapitulated up-regulation of major co-receptor genes associated with T cell dysfunction [36] — TIGIT, HAVCR2/TIM-3, PDCD1, and LAG3 (Fig. S13, Table S6). Known gene sets in T cell dysfunction [37, 36, 38] were significantly enriched in our up- *v.s*. down-regulated genes. Normalisr recovered biologically relevant and validated DE from human melanoma MARS-seq data.

We integrated the single-cell DE analysis onto the co-expression network of dysfunctional T cells to understand gene co-expression and clustering patterns from cell-to-cell variations (Fig. 4c, Table S7). We observed two distinct gene clusters of cell cycle (whose secondary PCs remained evident in co-expression despite the removal of its top PC) and type I interferon response (Table S8). We annotated the remaining, interconnected genes according to their known roles in T cell function in knowledge-base [39] and literature into one of seven categories: cytokines (and receptors), cytoskeleton, cytotoxic, MHC class I, MHC class II, T cell receptors (TCR), and co-receptors. Genes in the same functional category formed obvious regional co-expression clusters. Between the clusters, MHC class I genes neighbored MHC class II and TCR genes, whereas effector genes, including co-receptor, cytokine, and cytotoxic genes, were more closely linked and collectively up-regulated. CCL5, CSF1, CXCR6, and IL32 formed a separate co-expression cluster that is negatively associated with the rest of the cytokine program. Expression of dysfunctional genes, including CTLA4, LAG3, and TIGIT, were also correlated with cytokine activity. This cluster comprised a mixture of immune activation and exhaustion genes as well as sub-clusters divided by negative correlations. The co-expression network recovered by Normalisr suggests potential functional diversification in the dysfunctional T cell population, in agreement with previous discoveries in this field [40].

We then evaluated the functional associations recoverable from single-cell co-expression based on the over-abundance of co-expression edges between genes in each GO or KEGG annotation. Despite the previously reported difficulty in recapitulating GO annotations from single-cell co-expression networks from over 1,000 cells [41], we found that genes in 33 GO and 8 KEGG pathways had significantly more co-expressions than randomly assigned annotations (Bonferroni *P* ≤ 0.05, Table S9). These pathways encompassed a wide range of cell-type-specific and generic functions (Fig. 4d, Table S7). Overall, this analysis demonstrated that Normalisr recovered gene-level cellular pathways and functional modules in high-quality single-cell transcriptome-wide co-expression networks.

## Discussion

The current rise in scRNA-seq data generation represents a tremendous opportunity to understand gene regulation at the single-cell level, such as through pooled screens or co-expression. Here, we describe Normalisr as a unified normalization-association two-step inferential framework across multiple experimental conditions and scRNA-seq protocols. Normalisr efficiently infers gene regulations from pooled CRISPRi scRNA-seq screens and reduces false positives from gRNA cross-associations or off-target effects. Normalisr removes house-keeping modes in scRNA-seq data and infers transcriptome-wide, FDR-controlled co-expression networks that are consisted of cell-type-specific functional modules.

Normalisr addresses the sparsity and technical variation challenges of scRNA-seq with posterior mRNA abundances, nonlinear cellular summary covariates, and variance normalization. It fits in the framework of linear models [42] and achieves high performance over a diverse range of frequentist inference scenarios. Normalisr enables high-quality gene regulation and co-regulation analyses at the single-cell, single-gene level and at scale. However, Normalisr is not designed for nonlinear associations [43], causal regulatory networks through v-structures or regularized regression [44], or Granger causality with longitudinal information [4]. Integrative studies spanning multiple datasets may also require additional consideration for batch effects [45].

We established a restricted forward stepwise selection process to systematically dissect nonlinear effects of known confounders at a reduced burden of sensitivity loss or overfitting. This offers an unbiased and generalizeable approach to reduce problem-specific designs in normalization or association testing methods. We avoided informative prior or imputation to maintain independence between genes or cells for statistical inference, but nevertheless could outperform methods based on such techniques. Consequently, the unbiasedly determined nonlinear covariates were consistent across different scRNA-seq protocols such as 10X Genomics, MARS-seq, and CROP-seq at different sequencing depths.

Linear models possess immense capacities and flexibilities for scRNA-seq, such as ‘soft’ groupings for DE and kinship-aware population studies. At the same sequencing depth as bulk RNA-seq, scRNA-seq additionally partitions the reads between cells and cell types. This extra information provides substantial statistical gain and cell-type stratification.

## Methods

### Normalisr

#### Bayesian logCPM

Normalisr regards sequencing as a binomial sampling in the pool of mRNAs, *i.e*. the read/UMI count matrix 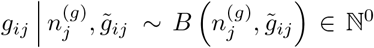 for gene *i* = 1,…, *n_g_* in cell *j* = 1,…, *n_c_*, where 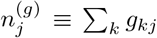 is the empirical sequenced mRNA count for cell *j* and 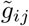 is the biological proportion of mRNAs from gene *i* in cell *j*. With a non-informative prior, the posterior distribution is 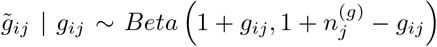. The minimum mean square error (MMSE) estimator for logCPM (here named Bayesian logCPM) is 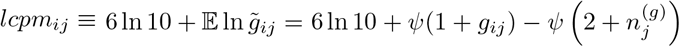, where *ψ* is the digamma function. Natural log is used throughout this paper.

#### Cellular summary covariates

To minimize spurious, technical gene inter-dependencies which may interfere with true co-expression patterns, we hypothesized and restricted covariate candidates within cellular summary statistics, including log total read count and the number of 0-read genes. To account for their potential nonlinear confounding effects, we used restricted forward stepwise selection with the optimization goal of uniformly distributed (linear) co-expression P-values on null datasets, *i.e*. simulated read count matrices based on real datasets but without any underlying co-expression.

Consider gene expression matrix 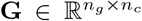 as Bayesian logCPM, and cellular summary covariates 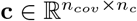 where *n_cov_* is the number of cellular summary covariates. Assume that **G** is linearly confounded at mean and variance levels by unknown, fixed, nonlinear column-wise functions 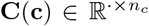, leading to false positives of gene co-expression even under the null hypothesis. We performed a Taylor expansion of each *C_i_* as

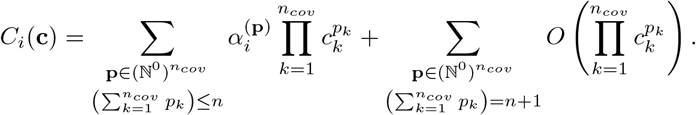

Therefore, nonlinear confounders **C** can be accounted for altogether with higher order sums of the original covariates **c**, provided the series converge quickly (subjecting to validation for the data type).

We iteratively included nonlinear covariates (parameterized by 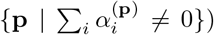. Starting from the constant intercept covariate set 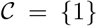, in each step we introduced the optimal candidate covariate to improve null co-expression P-value distribution from the set 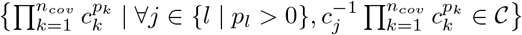. Note the restriction reduces the number of models/covariate candidates to a finite (and small) number, while only assuming that the aggregated effect of all *C_i_* functions is generically nonlinear, *i.e*. not very close to even or odd. The iteration ends when none could provide obvious improvement. The restricted forward stepwise selection avoids unnecessary covariates which degrade statistical power and reduce generalizability.

For Normalisr, the forward selection introduced three covariates: log total read count, its square, and the number of 0-read genes.

#### Existing covariate aggregation

Existing covariates can contain batches, pathways that are not the analytical focus (see GO pathway removal in co-expression networks), constant intercept term, etc. Aggregation of existing and cellular summary covariates is a simple concatenation followed by an orthonormal transformation (excluding intercept term).

#### Cell variance estimation

We modeled **lcpm** = *α*^(lcpm)^**C** +*ε*^(lcpm)^ in matrix form, where 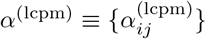 is the matrix of covariate *j*’s effect on gene *i*’s mean expression and is unbiasedly estimated with the maximum likelihood estimator (MLE) from linear regression as 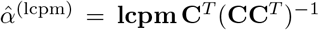. Then, we modeled the cell-specific error variance 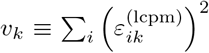 for cell *k* with ln **v** = *α*^(v)^**C** + *ε*^(*v*)^ with *ε*^(v)^ ~ *i.i.d* **N**(0, *σ*^2^). Its MLE as the estimated cell variance is 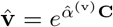 where 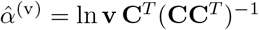. Note that maximum likelihood optimization with 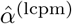 and 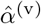 together fails due to overfitting and prioritization of few cells. This regression-based cell variance estimation retains changes of expression variance that are due to biological sources.

#### Gene expression variance normalization

Gene expressions were transformed to 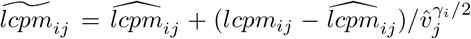 to normalize variance in the second term, where 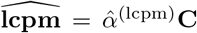. The scaling factor 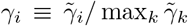, with 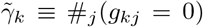, smoothly scales variance normalization effect to zero on highly expressed genes as their expressions are already accurately measured in scRNA-seq. Covariates’ variances were also scaled in full to 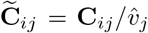 accordingly with the exception of categorical covariates left unchanged in one-hot encoding.

#### Gene differential and co-expression hypothesis tests

Using normalized expression and covariates, differential expression was tested with linear model 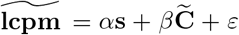, where *s* = 0,1 indicates cell set membership (case *v.s*. control) and *ε* ~ *i.i.d N*(0, *σ*^2^). The log fold-change was estimated as 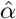 using maximum likelihood. The two-sided P-value was computed for the null hypothesis *α* = 0 using the exact null distribution of the proportion of explained variance (by **s**), as 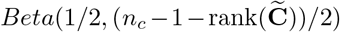. Co-expression between genes i,j was tested with 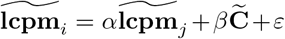, with the same setup otherwise. Their (conditional) Pearson correlation and P-value were computed from *α*, and are symmetric between *i* and *j*.

#### Outlier cell removal

For outlier removal, inverse of (estimated) cell variance was modeled with normal distribution. Given the prior bound of outlier proportion as *r*, the iterative outlier detection method started with *r* to 1 – *r* percentiles as non-outlier cells. In each iteration, a normal distribution was fitted with MLE for the inverse variances of non-outliers, and was used for outlier test of all cells with two-sided P-values. Cells below the given Bonferroni adjusted P-value threshold were regarded as outliers in the next iteration. This was repeated until convergence and the prior bound of outlier proportion *r* was checked. The removal process takes the prior bound of outlier proportion *r* and the P-value threshold as parameters.

#### GO pathway removal in co-expression networks

The GO pathway removal took an iterative process given the significance cutoff for co-expression Q-value. In each iteration, Normalisr first identified the top 100 principal genes in the co-expression network, defined as those with most co-expressed genes (passing the significance threshold). To find the dominating pathway for co-expression, Normalisr performed GO enrichment analysis on the these regulators using all genes after quality control as background. The most enriched GO term (by P-value) was manually examined for contextual relevance and house-keeping role. The researcher then chose to proceed the removal or stop iteration.

If removal was elected, Normalisr first removed existing covariates linearly from the (Normalisr normalized) transcriptomic profile. Then Normalisr extracted the top principal component of the subset transcriptomic profile of genes in the GO term. This principal component was added as an extra covariate, before producing a new co-expression network and repeating the iteration.

This process was performed iteratively until the most enriched GO term was contextually relevant, as judged by the researcher. The only manual input of the iterative process was when to stop iteration.

### Method evaluations

#### Random partition for null DE

Homogenous cells from the same dataset, cell-type, and condition were partitioned to two sets randomly for 100 times for null differential expression. Each random partition had a different partition rate, sampled uniformly between 5% and 50%.

#### Existing normalization and imputation methods

We used the default configurations in sctransform (variance stabilizing transformation, 0.2.1), bayNorm (separately with mode_version or mean_version, 1.2.0), Sanity (1.0), DCA (0.2.3), and MAGIC (1.2.1), with 50 latent dimensions for MAGIC. Their differential and co-expression used the same linear model but without cellular summary covariates. Sctransform’s log10 expression was converted to natural log for logFC comparison. ScImpute [46] and ZINB-WaVE [47] were not included because they could not accommodate the data size with excessive memory and/or time requirements. ScVI [48] could not fit into our comparison because it did not recommend P-value based FDR estimation.

#### Existing DE methods

For edgeR and MAST, we exactly replicated the codes in [8], using edgeRQLFDetRate for edgeR (3.26.8) [9] and MASTcpmDetRate for MAST (1.10.0) [1]. EdgeR’s log2 fold change was converted to natural log for comparison.

#### Evaluation of null P-value distribution in hypothesis testing

For differential gene expression, we split genes to 10 equally sized groups by number of expressed cells, low to high. We then evaluated the null distribution of P-values for each group separately. This can reveal null P-value biases that depend on expression level. For the same purpose, we evaluated the null distribution of gene co-expression P-values for each pair of gene groups separately.

Two-sided KS test was performed on each of the 10 groups of null P-values for DE, or 55 groups for co-expression, to evaluate the how they represent a standard uniform distribution. The resulting 10 or 55 KS test P-values were drawn and compared across different methods with violin plot. Larger KS test P-values indicate better recovery of uniform null P-values.

#### Running time evaluation

Each method was timed after quality control on a high-performance computer of 64 CPU cores excluding disk read/write. Sanity can only use file input/output which incurs a minor timing overhead in our evaluation pipeline.

#### Synthetic dataset of null co-expression

To produce a synthetic read count matrix 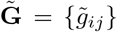 without co-expression of *ñ_c_* cells and *ñ_g_* genes, yet mimicking a real, pre-quality control dataset with read count matrix **G** = {*g_ij_*}, of *n_c_* cells and *n_g_* genes, we took the following steps.

1. Compute empirical distributions of read counts per cell 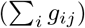 as 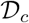, and total read proportions per gene 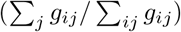 as 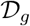.
2. *n_j_*: Sample/Simulate each cell *j*’s total read count from 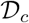, and scale them by *ñ_g_/n_g_*.
3. *p_i_*: Sample each gene *i*’s mean read proportion from 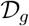.
4. *b_ij_ ~ i.i.d N*(0, *σ*^2^): Simulate biological variations.
5. 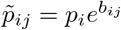: Compute expression proportion.
6. 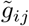: For cell *j*, sample *n_j_* reads for all genes, with weight 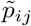 for gene *i*.

The simulation contains 3 parameters: the number of cells *ñ_c_*, the number of genes *ñ_g_*, and biological variation strength *σ*. It reflects the confounding effect of sequencing depth in real datasets, which is lost by permutation-based null datasets. This model is similar with powsimR’s negative binomial model [49], except that it separates biological variations from the binomial detection process, allows for extra method evaluations with the ground-truth of biological mRNA proportions 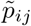, and accepts biological variation strength *σ* as input.

#### Differential expression logFC estimation evaluations

Estimated logFCs were compared against the known ground-truth in synthetic datasets for method evaluation. To decompose estimation errors into bias and variance, we trained a univariate, intercept-free linear model to predict estimated logFCs with the ground-truth logFCs. The resulting regression coefficient represents the overall bias in logFC scale estimation. The goodness of fit (Pearson *R*^2^) indicates the variance of logFC estimation. This evaluation was performed on separate gene groups from low to high expression levels (in terms of proportion of expressed cells) to better characterize method performances. The ideal method should provide low bias, low variance, and stable bias that changes minimally with expression level.

#### Reproducibility across scRNA-seq protocols and experimental conditions

We used the MARS-seq dataset [26] of human melanoma tissue samples at a lower sequencing depth to evaluate Normalisr’s performances across different scRNA-seq protocols and experimental conditions. Only annotated non-live, dysfunctional T cells from frozen samples were selected as a relatively homogeneous population for method evaluation. As a secondary evaluation, we excluded time-consuming and sub-optimal DE methods edgeR and MAST. All compatible evaluations were repeated on the MARS-seq datasets.

### General methods

#### ScRNA-seq quality control (QC)

In this study, we restricted our analyses to cells with at least 500 reads and 100 genes with non-zero reads, and genes with non-zero reads both in at least 50 cells and in at least 2% of all cells for 10x technologies, or in at least 500 cells and in at least 2% of all cells for MARS-seq, due to their different read count distributions in this study. The QC was performed iteratively until no gene or cell is removed. Simulated datasets followed the criteria of the corresponding real dataset technology.

#### Gene Ontology

Gene Ontology enrichment with `GOATOOLS` (0.8.4) [50] was restricted to post-QC genes as background, except in co-expression network modules where all known genes were used. To avoid evaluation biases for our expression-based analyses, for all GO analyses we excluded GO evidences that may also have an expression-based origin (IEP, HEP, RCA, TAS, NAS, IC, ND, and IEA). All three GO categories were used. GO enrichment P-values are raw unless stated otherwise.

### High-MOI CRISPRi scRNA-seq analyses

#### Guide RNA cross-association test

Odds ratio of gRNA intersection was computed for every gRNA pair, with one-sided hypergeometric P-values.

#### Types of gRNA-gene pairs

- Naive/aware, target: NTC gRNA *v.s*. genes targeted by any gRNA at its TSS
- Naive/aware, other: NTC gRNA *v.s*. genes not targeted by any gRNA at its TSS
- TSS, target: TSS-targeting gRNA *v.s*. the gene it targets
- TSS, other: TSS-targeting gRNA *v.s*. genes it doesn’t target
- NTC: NTC gRNA *v.s*. any gene
- Enhancer: Enhancer-candidate-targeting gRNA *v.s*. any gene

#### Association P-value and logFC histograms

Histograms were constructed with 50 bins, whilst being symmetric for logFC. Separate histograms were drawn for different gRNA-gene pair types. Absolute histogram errors were estimated as 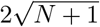 where *N* is the number of occurrences at each bin.

#### FPR and Q-value estimations

FPR was estimated with 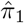 in Storey’s method [28] using `fdrtools` [51], separately for each gRNA-gene pair type. To account for variations in gRNA specificity and gene role, Q-values were estimated with Benjamini-Hochberg procedure, together for all gRNA-gene pairs in the ‘TSS, target’ type, and separately for each gRNA in other types.

#### Competition-aware method

The competition-aware method regards all other, untested gRNAs as covariates. For efficiency, these covariates were introduced at the time of association testing and were assumed to only affect the mean expression, and all covariates were reduced to top 500 PCs if more than 10,000. Other numbers of top PCs have been tried and found reliable against no dimension reduction on the small-scale dataset.

#### Comparison with existing CRISPR scRNA-seq screen analysis methods

We downloaded the author’s deposition of SCEPTRE’s gene-level P-values on the same full-scale K562 study for comparison of sensitivity on the repression effects of TSS-targeting gRNA as positive controls. To enable comparison with SCEPTRE, gRNA level P-values from Normalisr were combined to gene level with Fisher’s method. P-value strength comparison was limited to positive control genes that were significant according to either Normalisr or SCEPTRE (Bonferroni *P* < 0.05). P-values for non-cis-effects (including from NTCs) were not available from SCEPTRE for sensitivity or specificity comparison.

ScMAGECK was not compared for sensitivity because it did not finish computation within two weeks and scMAGECK’s P-values were not publicly available for the same study. Running time projections were based on [12] for SCEPTRE and Fig. 2c for edgeR and MAST.

#### CRISPRi off-target rate

For each gRNA-targeted gene, a weaker and a stronger gRNA targeted its TSS, according to the P-value of their linear associations. The raw off-target rate was defined as the proportion of the gRNA-targeted gene’s inferred targets (*Q* ≤ 0.05 or 10^-5^) based on the weaker gRNA that were not reproducible with the stronger gRNA (*P* ≥ 0.1). It is also the FDR when evaluating the weaker gRNA’s associations using the stronger one’s as gold standard. The estimated off-target rate is min(raw off-target rate/(1 – 0.1), 1) to extrapolate and account for the proportion of insignificant associations with the stronger gRNA that was lost in choosing *P* ≥ 0.1.

#### Inferring gene regulations from trans-associations

Inferred gene regulations must satisfy all the following conditions for both gRNAs in both screens.

- Significant repression of targeted gene: *Q* ≤ 0.05 and logFC ≤ –0.2.
- Significant association with trans-gene: *Q* ≤ 0.2, |logFC| ≥ 0.05, relative (to logFC of targeted gene) |logFC| ≥ 0.05.
- No other gene repressed within 1Mbp up- and down-stream window of targeted TSS: *P* ≤ 0.001 and logFC ≤ –0.1.

### Dysfunctional T cells in human melanoma

#### Dataset

We downloaded the read count matrix and meta-data from GEO, after QC from the original authors. We regarded the following as categorical covariates that may confound gene (co-)expression: batches from amplification, plate, and sequencing, as well as variations from patient, cell alive/dead, sample location, and sampling processing (fresh or frozen). In our QC, cells from donors that had fewer than 5 cells of the same cell type were discarded. With the prior bound for outlier proportion as 2%, low-variance outlier cells with Bonferroni *P* ≤ 10^-10^ were removed from downstream analyses with Normalisr. Pseudo-genes and spike-ins were treated indifferently in statistical inference.

#### Dysfunctional gene overlap testing

Overlap testing was performed between known dysfunctional genes (from literature) and genes up-regulated (*Q* ≤ 0.05) in dysfunctional *v.s*. naive T cells. Background gene set of this dataset was selected as up- or down-regulated genes (*Q* ≤ 0.05). Only known dysfunctional genes found in the background set were considered for hypergeometric testing P-value.

#### Co-expression network

We computed Q-value networks from raw P-values separately for each gene against all other genes, to account for different gene roles such as principal genes or master regulators [21]. For co-expression network in dysfunctional T cells, we used a strong cutoff (*Q* ≤ 10^-15^) to prioritize more direct gene interactions. The final co-expression network focused on the major connected component of the full co-expression network after two iterations of GO pathway removal. Within the final network, we also removed a relatively separate cluster consisted solely of non-coding mRNAs and spike-ins (Fig. S12).

#### Over-abundance of co-expressions within annotation group

We used `GOATOOLS` [50] and `BioServices` [52] for GO and KEGG pathways respectively. We restricted the gene sets to having at least 2 edges but at most half in the given co-expression network. To reduce multiple testing, for each GO term, its parent term is excluded if it has the same annotation on the network. P-values for edge over-abundance were computed with random annotation assignment with the same number of annotated nodes in the network.

## Availability of data and materials

Normalisr (0.6.0) is publicly available at https://github.com/lingfeiwang/normalisr. Published datasets were downloaded from Gene Expression Omnibus with accession numbers: GSM2406681 (Perturb-seq), GSE123139 (MARS-seq), GSE99254 (Smart-Seq2), GSE120861 (CROP-seq), GSM2138664 (shRNA DDX3X knock-down, ENCODE identifier: ENCSR000KYM), and GSM2138876 (shRNA control, ENCODE identifier: ENCSR913CAE).

## Acknowledgements

LW would like to thank Daniel Graham, Kirk Gosik, and Heping Xu for early discussions, Jacques Deguine for insights in T cell biology, Ramnik Xavier for funding support, and Jacques Deguine and Qian Qin for extensive feedback on this manuscript. LW would like to acknowledge the ENCODE Consortium and Brenton Graveley’s Lab for generating the shRNA DDX3X knock-down datasets. LW personally appreciates Lynn Rekhi’s administrative assistance during the COVID-19 outbreak.

---

**Supplementary Figure 1:**
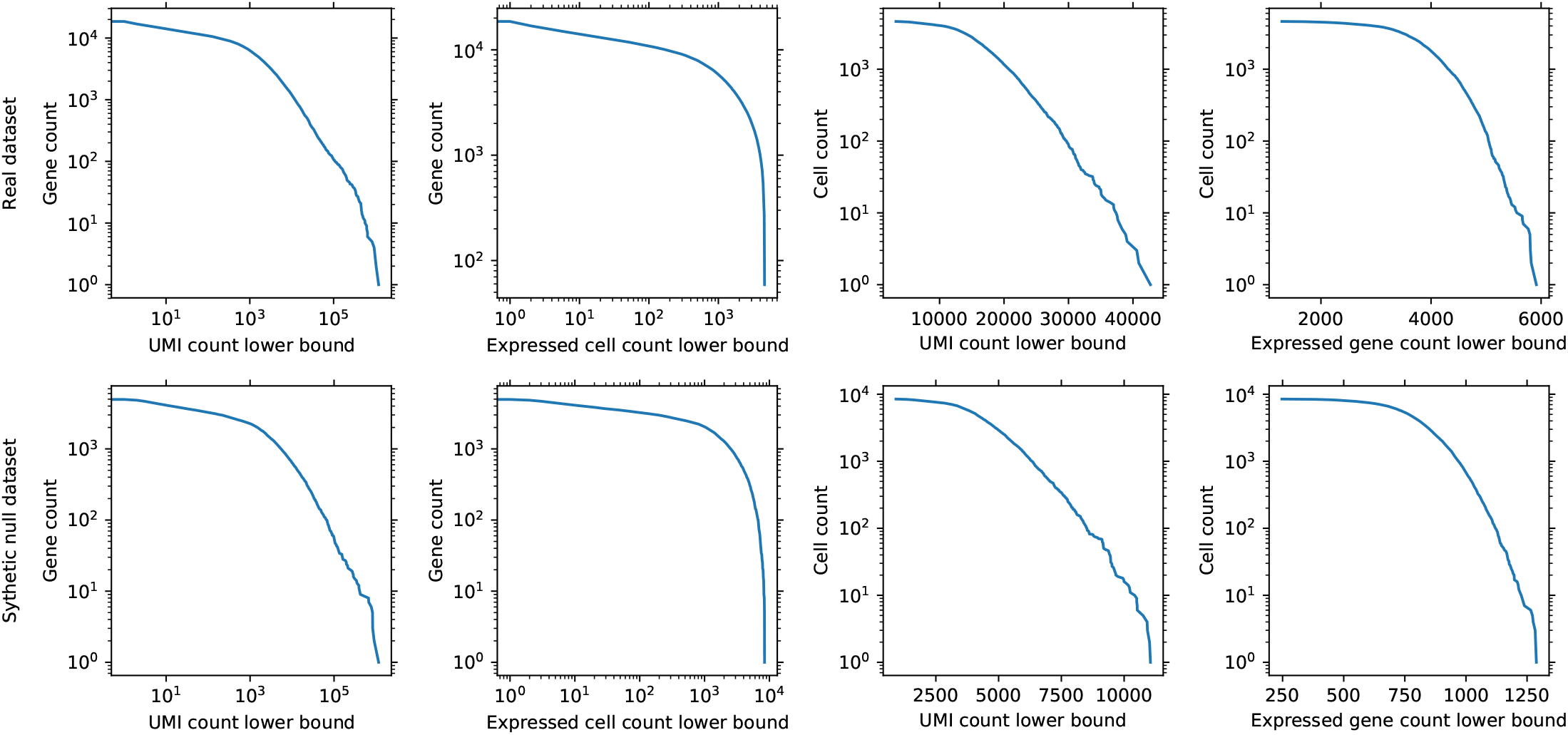
Synthetic null co-expression dataset (5,017 genes in 8,472 cells pre-QC and 3,036 genes in 8,472 cells post-QC) mimicked the read count distributions of the real dataset (18,583 genes in 4,622 cells) in terms of survival functions. Expressed cell for a given gene (x axis in columns 2 and 4) is defined as those cells with non-zero read count for the gene.

**Supplementary Figure 2:**
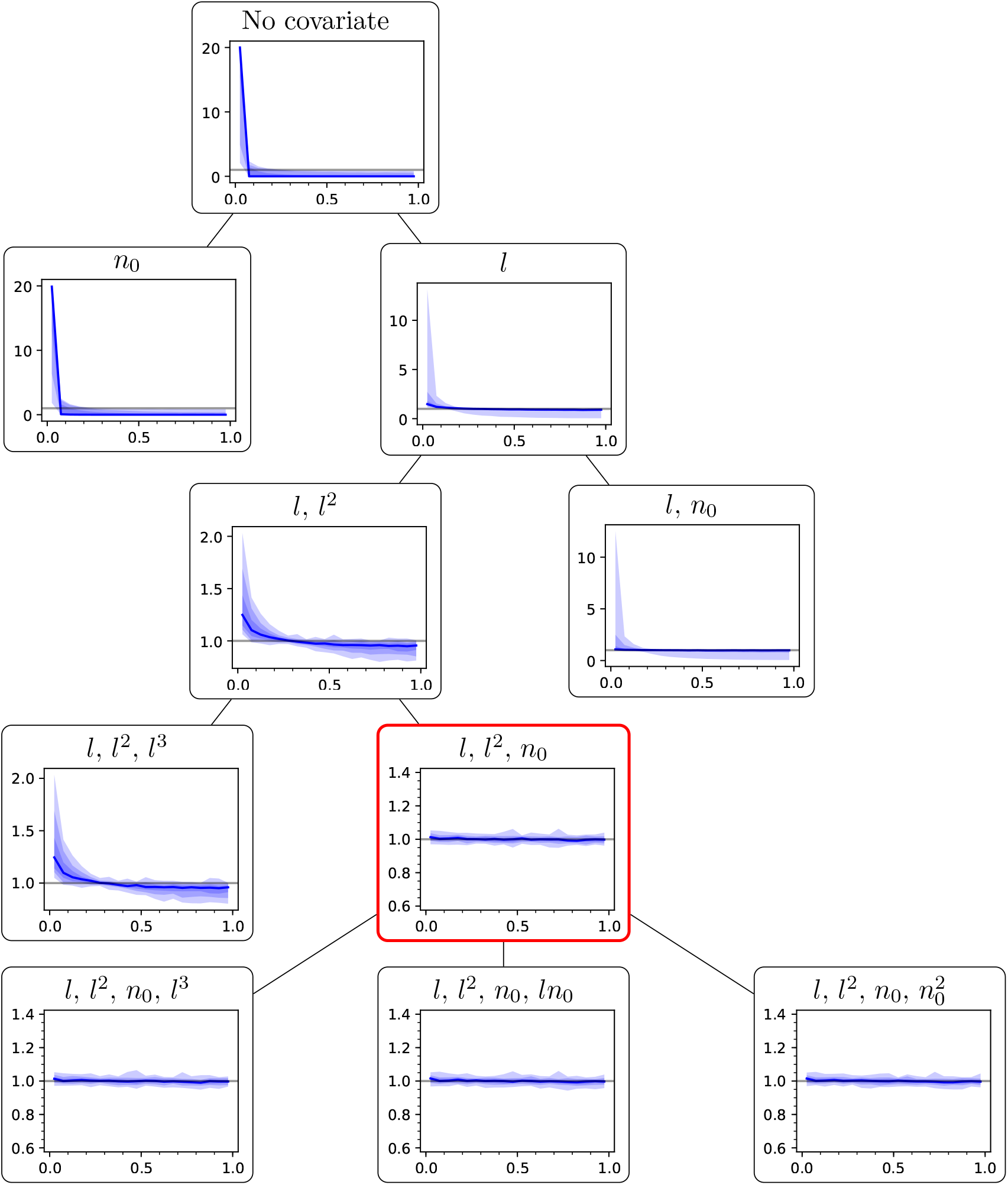
Decision tree to introduce covariates iteratively as a Taylor expansion. Each step (top to bottom) shows the covariates (*l*: total log read count in each cell, *n*_0_: number of 0-read genes in each cell, besides constant intercept) and the resulting histograms (Y) of null co-expression P-values (X), based on the same dataset and drawing style as in Fig. 2c. The covariate set was optimized towards a uniform distribution of null co-expression P-values. Red: the covariate set with the best P-value histogram and fewest covariates was selected for Normalisr.

**Supplementary Figure 3:**
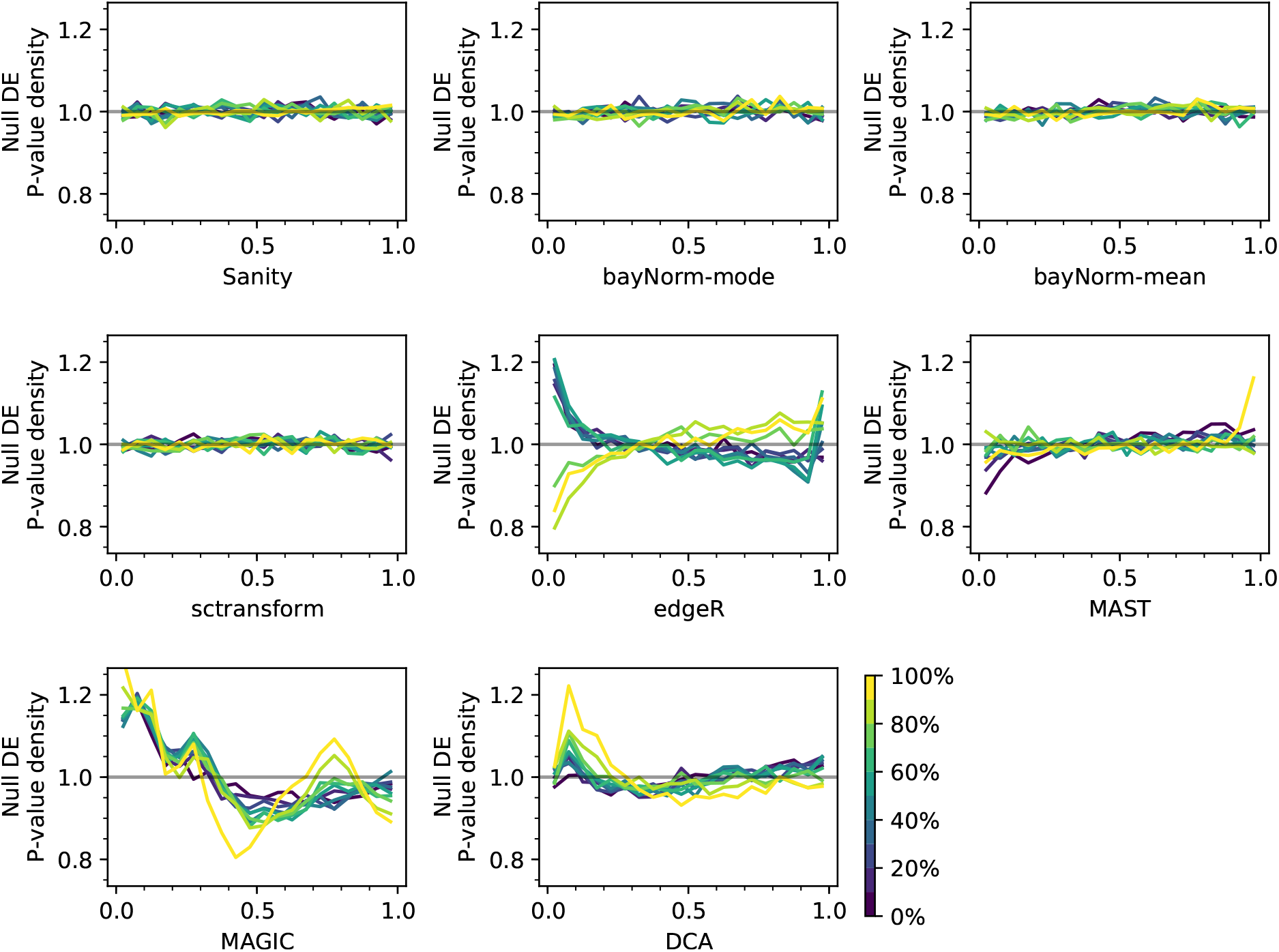
Method performances vary in recovering uniformly distributed null P-values in single-cell DE (X) as shown by histogram density (Y). Genes were evaluated in 10 bins from low to high expression (color).

**Supplementary Figure 4:**
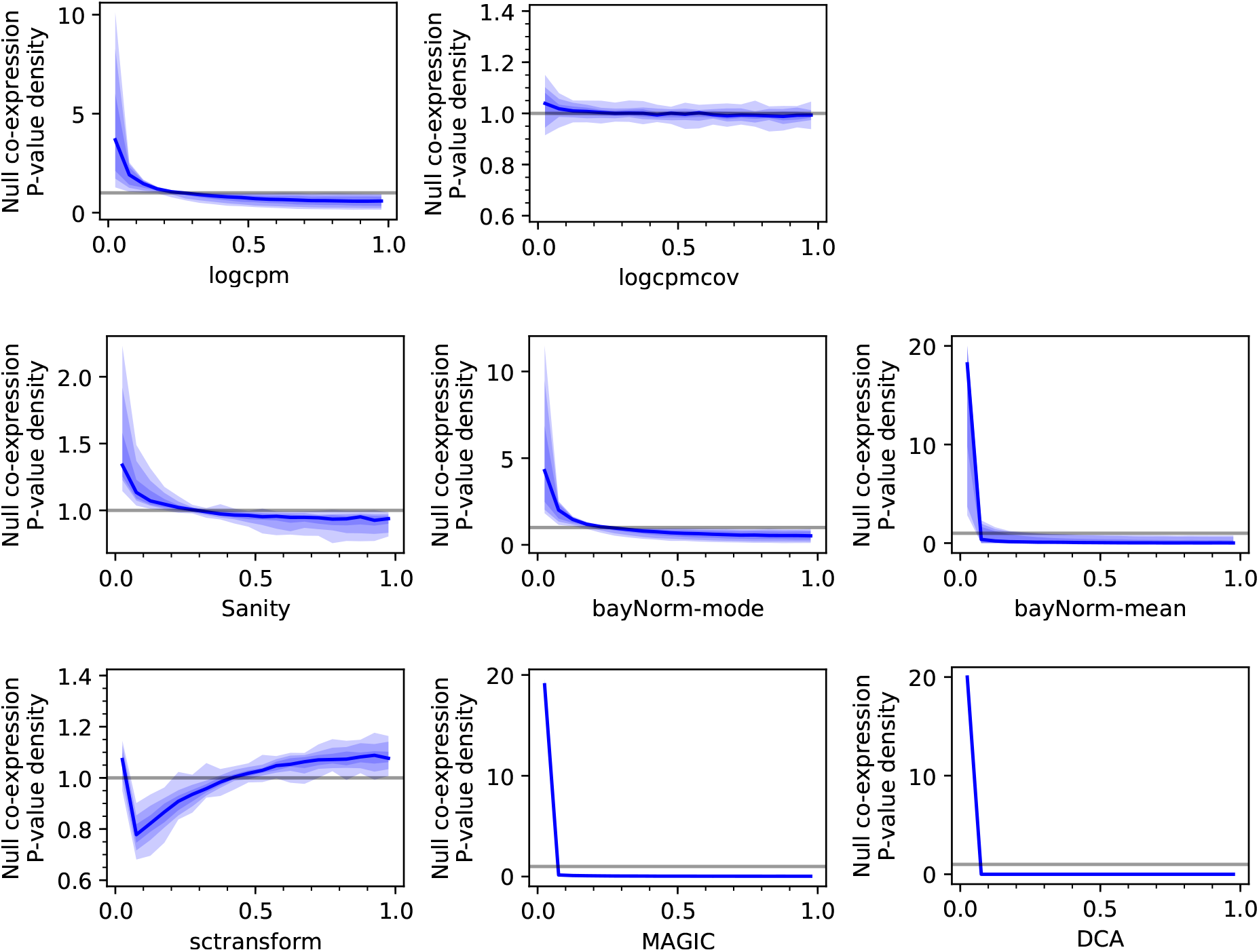
Other methods recovered distorted null P-value of single-cell co-expression (X) as shown by histogram density (Y). Left to right and top to bottom: log(CPM+1), log(CPM+1) with cellular summary covariates, Sanity, bayNorm-mode, bayNorm-mean, sctransform, MAGIC, and DCA. Genes were split into 10 equal bins from low to high expression. The null P-value distribution of co-expression between each bin pair formed a separate histogram curve. Central curve shows the median of all histogram curves. Shades show 50%, 80%, and 100% of all histograms. Gray line indicates uniform distribution.

**Supplementary Figure 5:**
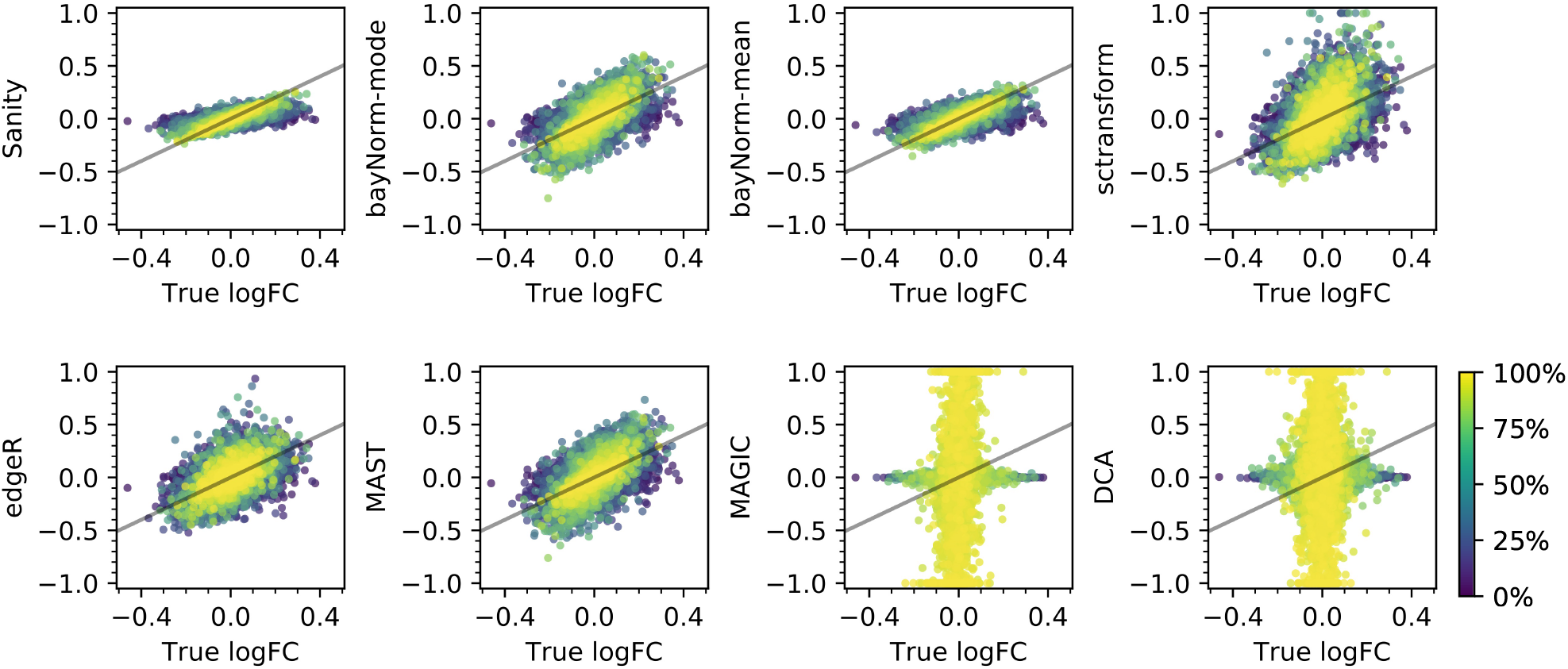
Other methods had stronger bias and/or variance in logFC estimation from synthetic data. Recovered logFCs (Y) were compared against the synthetic ground-truth (X) for genes from low to high expression (color) for different methods. Gray lines indicate X=Y.

**Supplementary Figure 6:**
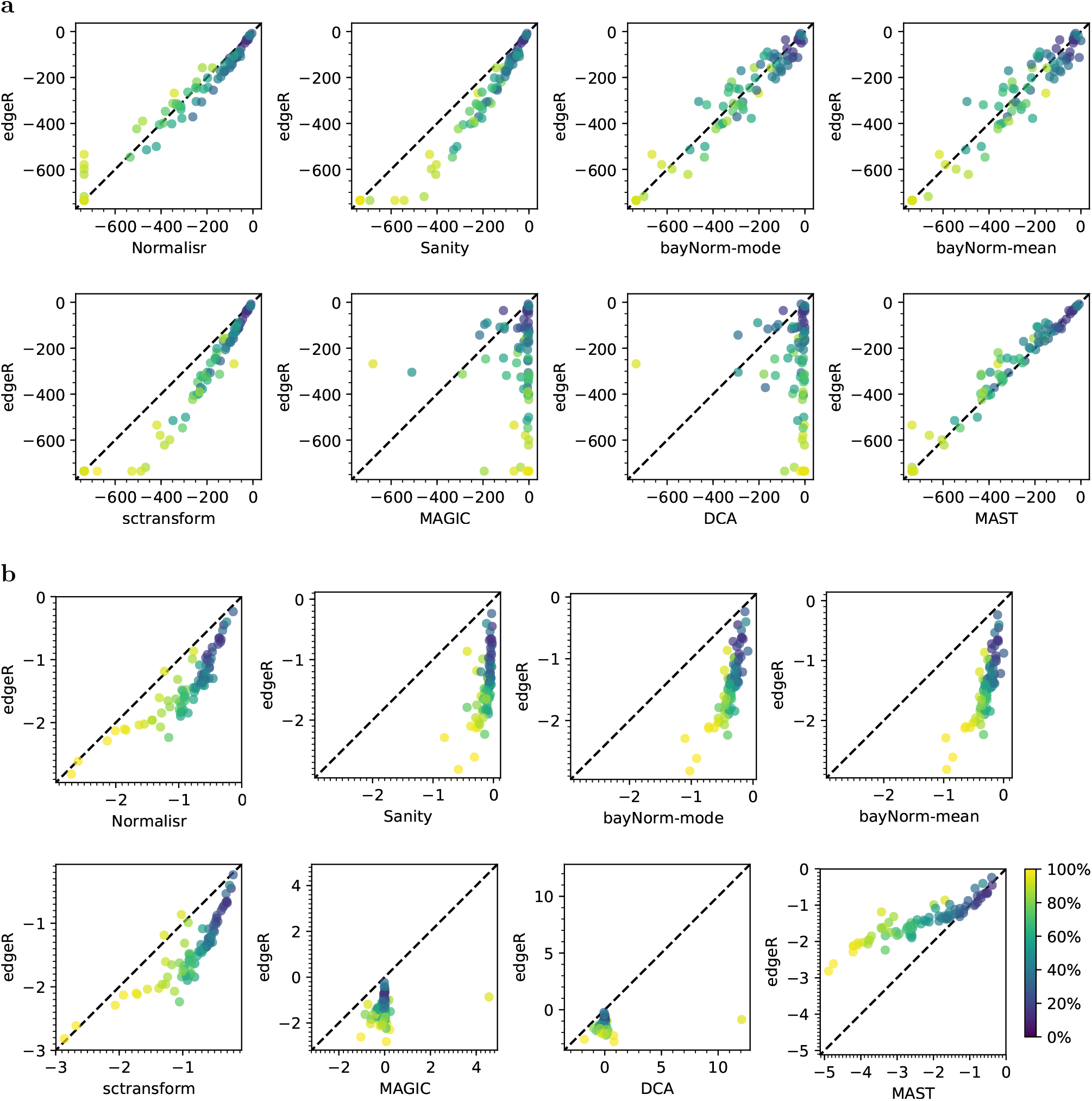
Normalisr obtained high sensitivity and accuracy in DE among CRISPRi positive controls. DE log P-value (**a**) and logFC (**b**) of the same CRISPRi gRNA-target pairs are compared between edgeR (Y) and each other method (X). Genes are colored according to the proportion of expressed cells. Dashed line indicates X=Y.

**Supplementary Figure 7:**
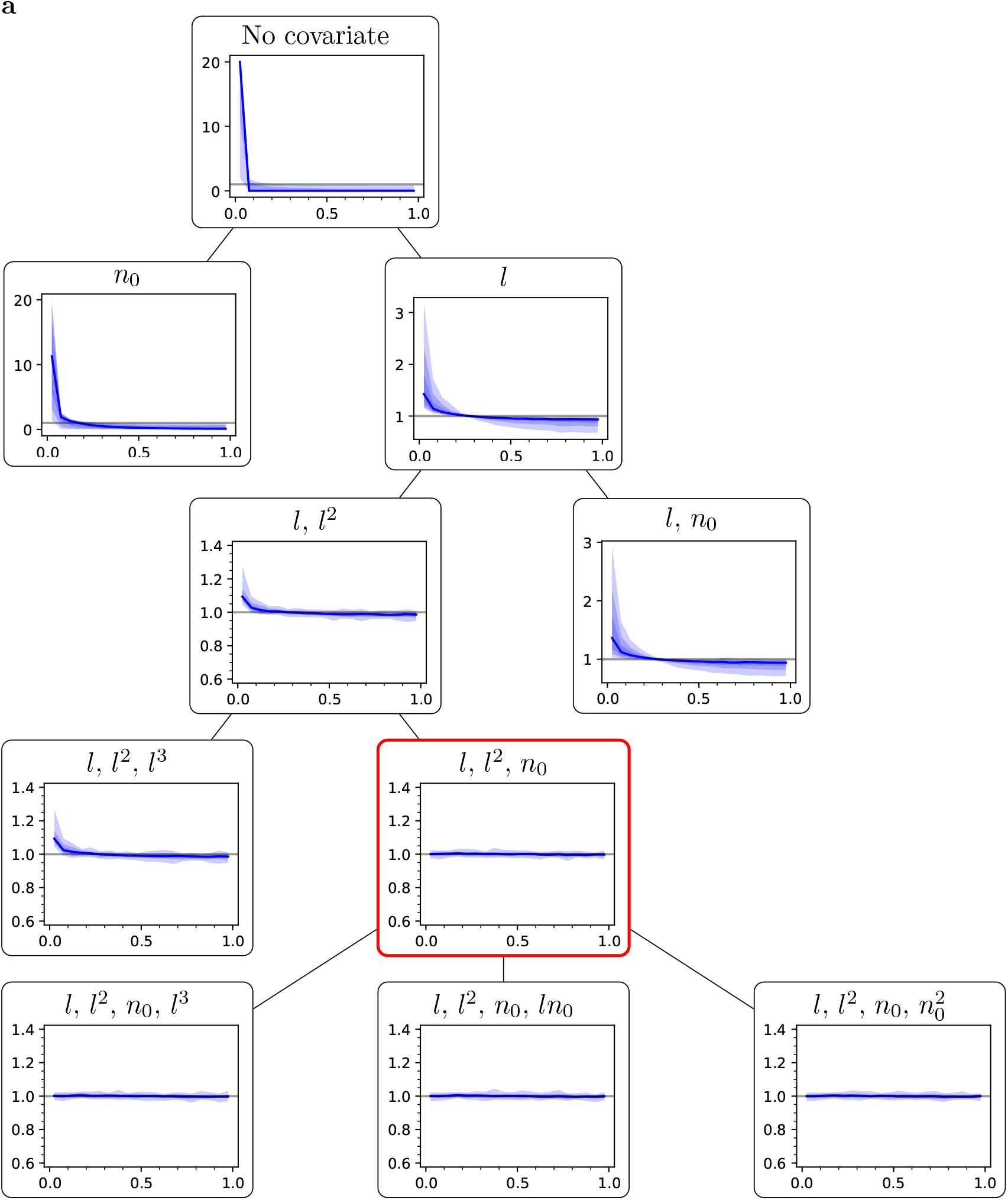
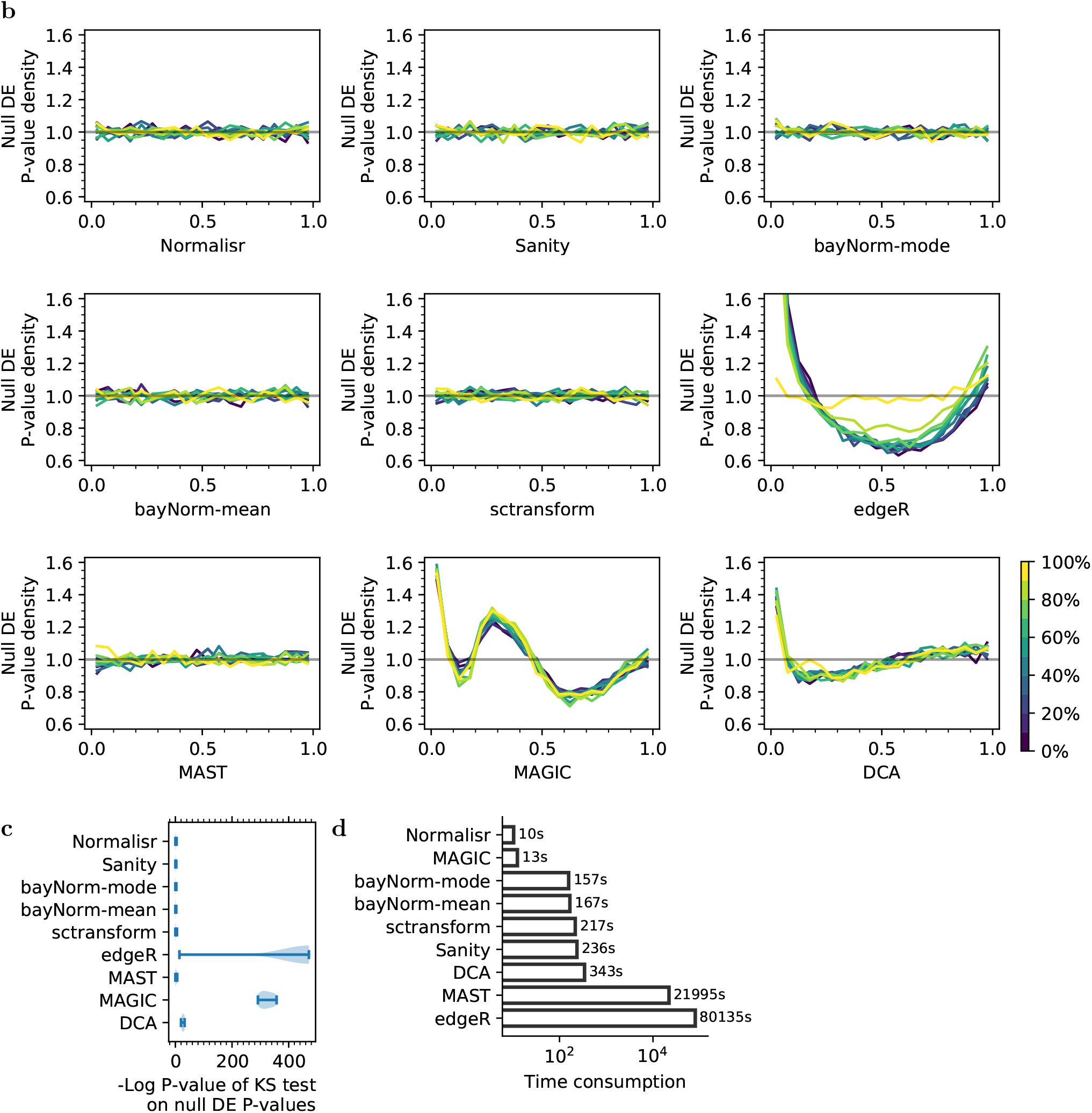
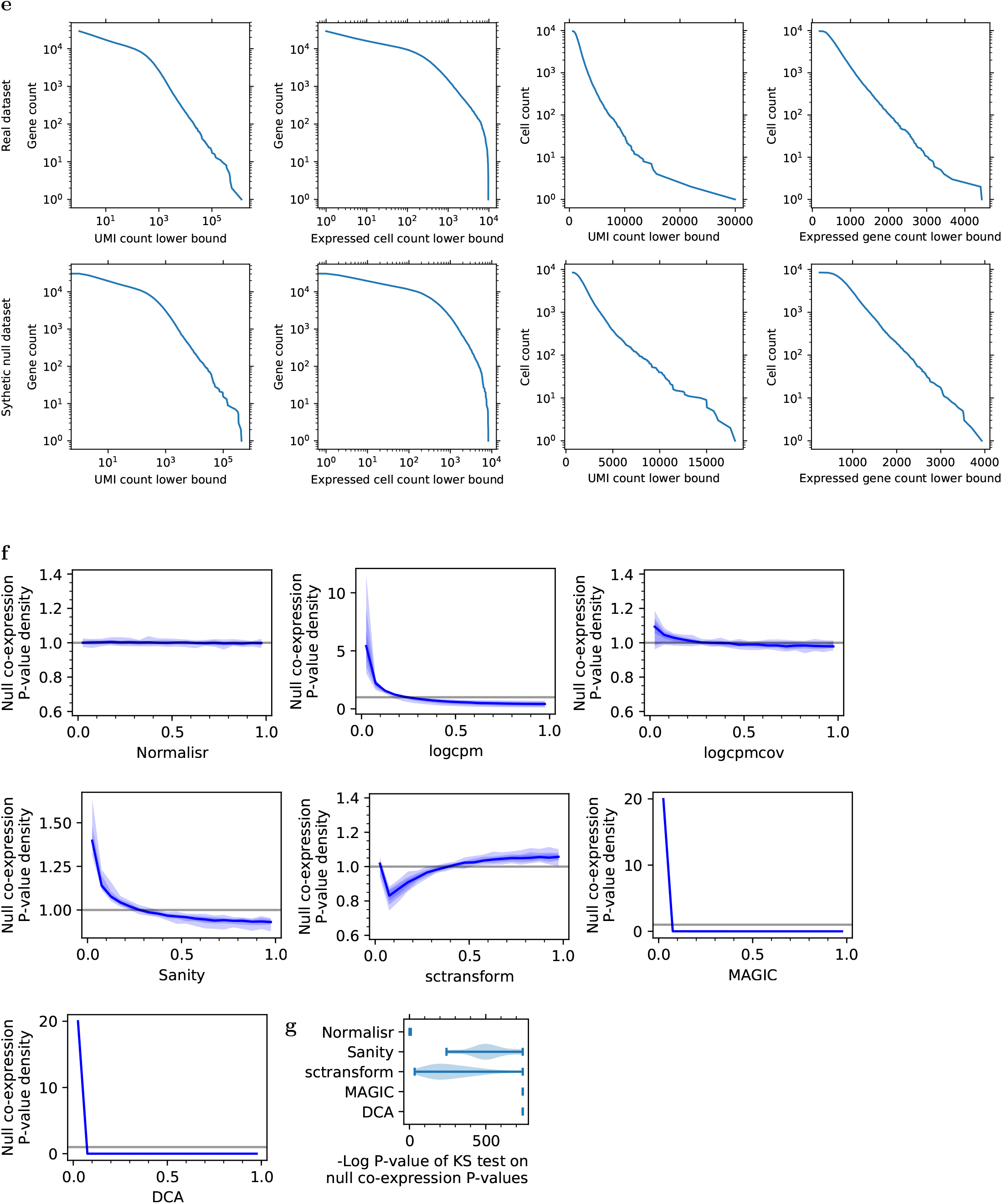
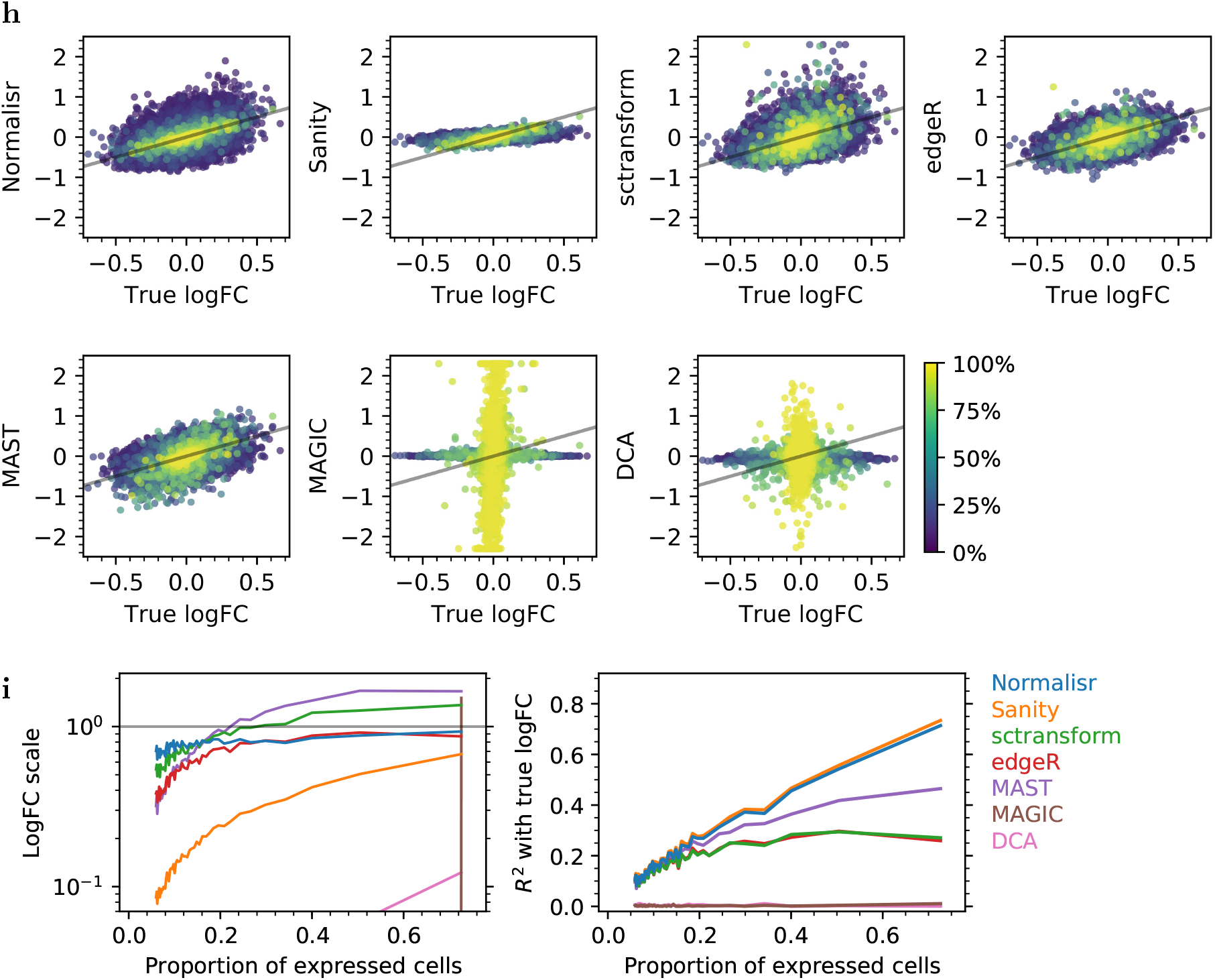
Evaluation results were reproducible at a lower sequencing depth on the MARS-seq dataset of dysfunctional T cells from frozen human melanoma tissue. Evaluations were reproduced for Fig. S2 (in panel **a**), Fig. 2a and Fig. S3 (**b**), Fig. 2b (**c**), Fig. 2c (**d**), Fig. S1 (**e**), Fig. 2d and Fig. S4 (**f**), Fig. 2e (**g**), Fig. 2f and Fig. S5 (**h**), and Fig. 2g (**i**). **a** Choice of nonlinear cellular summary covariates with restricted forward stepwise selection remained consistent for Normalisr on the MARS-seq of frozen human melanoma tissues at a lower sequencing depth. **bc** Normalisr, Sanity, bayNorm, sctransform, and MAST had uniformly distributed null P-values (X) as shown by histogram density (Y) in single-cell DE. **d** Normalisr was much faster than other normalization methods. **e** Synthetic null co-expression dataset (32,854 genes in 8,472 cells pre-QC and 4,768 genes in 8,471 cells post-QC) mimicked the read count distributions of the real dataset (29,161 genes in 9,760 cells pre-QC) in terms of survival functions. **fg** Only Normalisr had uniformly distributed null P-values in single-cell co-expression (X) from synthetic data, as shown by histogram density (Y), that mimicked MARS-seq of dysfunctional T cells from frozen human melanoma tissues at a lower sequencing depth. **h** Normalisr accurately recovered logFCs (Y) when compared against the synthetic ground-truth (X) for genes from low to high expression (color). Gray line indicates X=Y. **i** Normalisr accurately recovered logFCs with low variance (right, Y as *R*^2^) and low bias (left, Y as regression coefficient; horizontal gray line indicates bias-free performance) when evaluated against synthetic ground-truth with a linear regression model separately for genes grouped from low to high expression (X) on logFC scatter plots (**h**). BayNorm failed parts of the evaluations with undocumented errors and could not be compared.

**Supplementary Figure 8:**
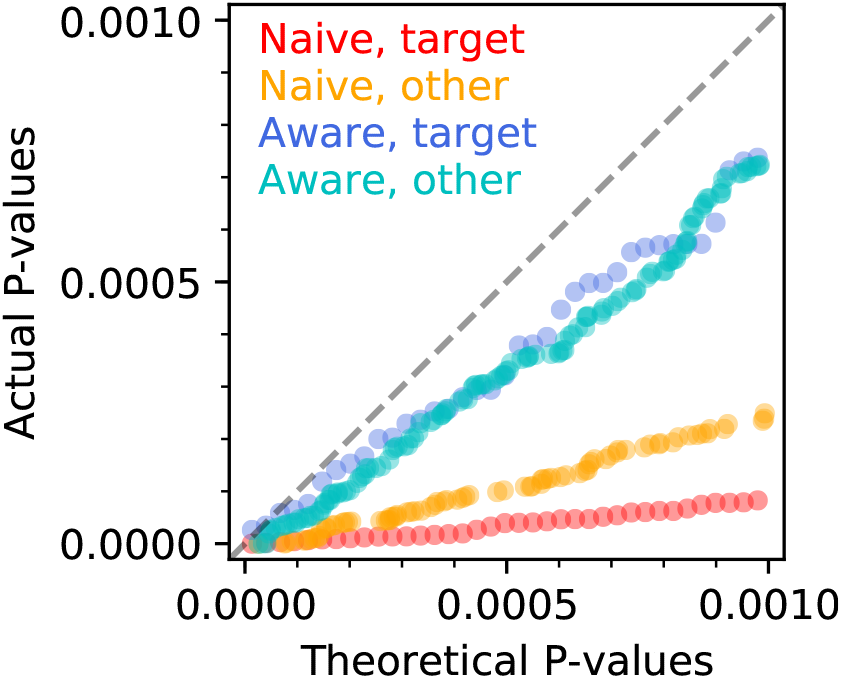
Quantile-quantile plot of theoretical (X) and actual (Y) P-values of CRISPRi DE from NTC gRNAs. DE was performed separately with competition-**naive** and -**aware** methods and separately for genes **target**ed by positive control gRNAs at the TSS and **other** genes. Data points were randomly down-sampled to avoid overly dense plots.

**Supplementary Figure 9:**
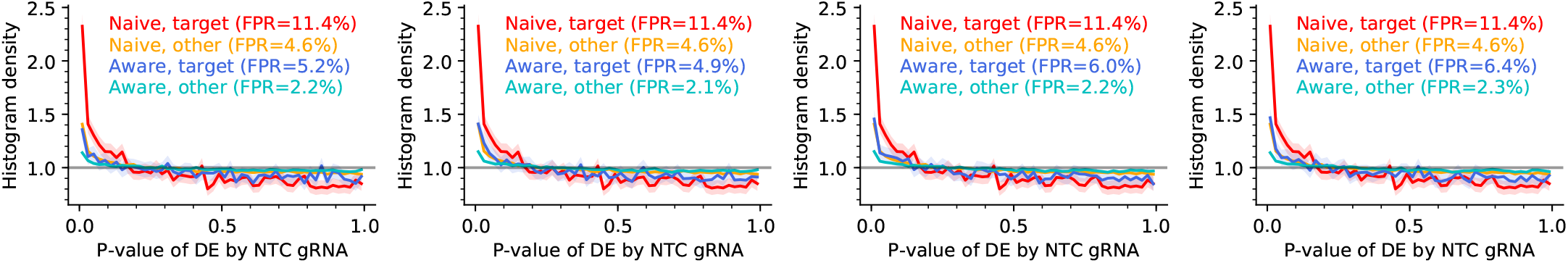
Top principal components of gRNA variation could account for most of the confounding effects on the small-scale CRISPRi screen. Density histograms (Y) of P-values (X) from the competition-aware method remained almost identical between considering (left-to-right) all other gRNAs and their top 500, 200, or 100 principal components.

**Supplementary Figure 10:**
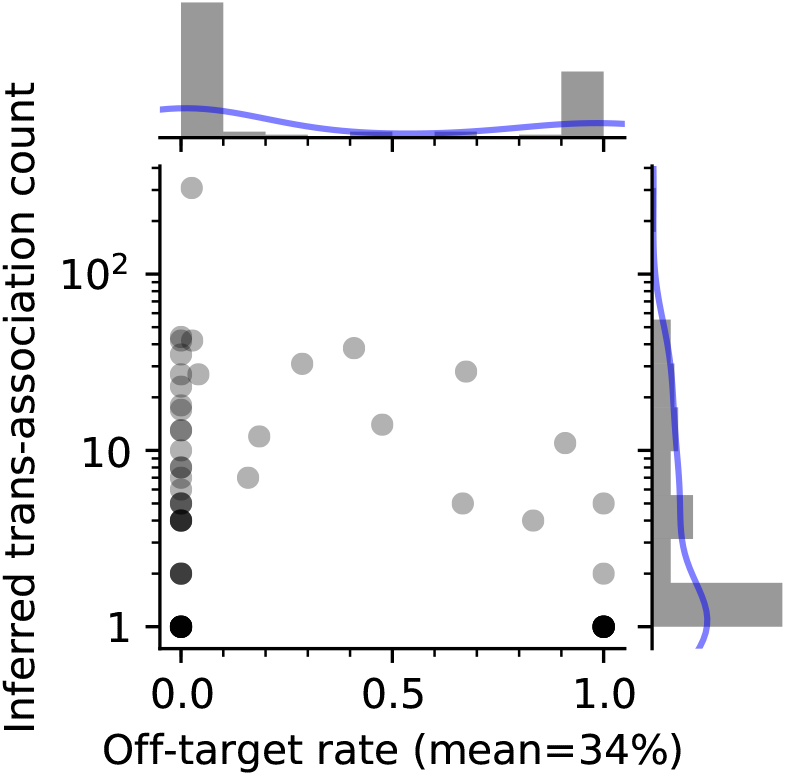
CRISPRi off-target effects were weaker than genuine trans-associations. Estimated mean off-target rate (X) reduced to 34% at *Q* ≤ 10^-5^ for significant trans-associations.

**Supplementary Figure 11:**
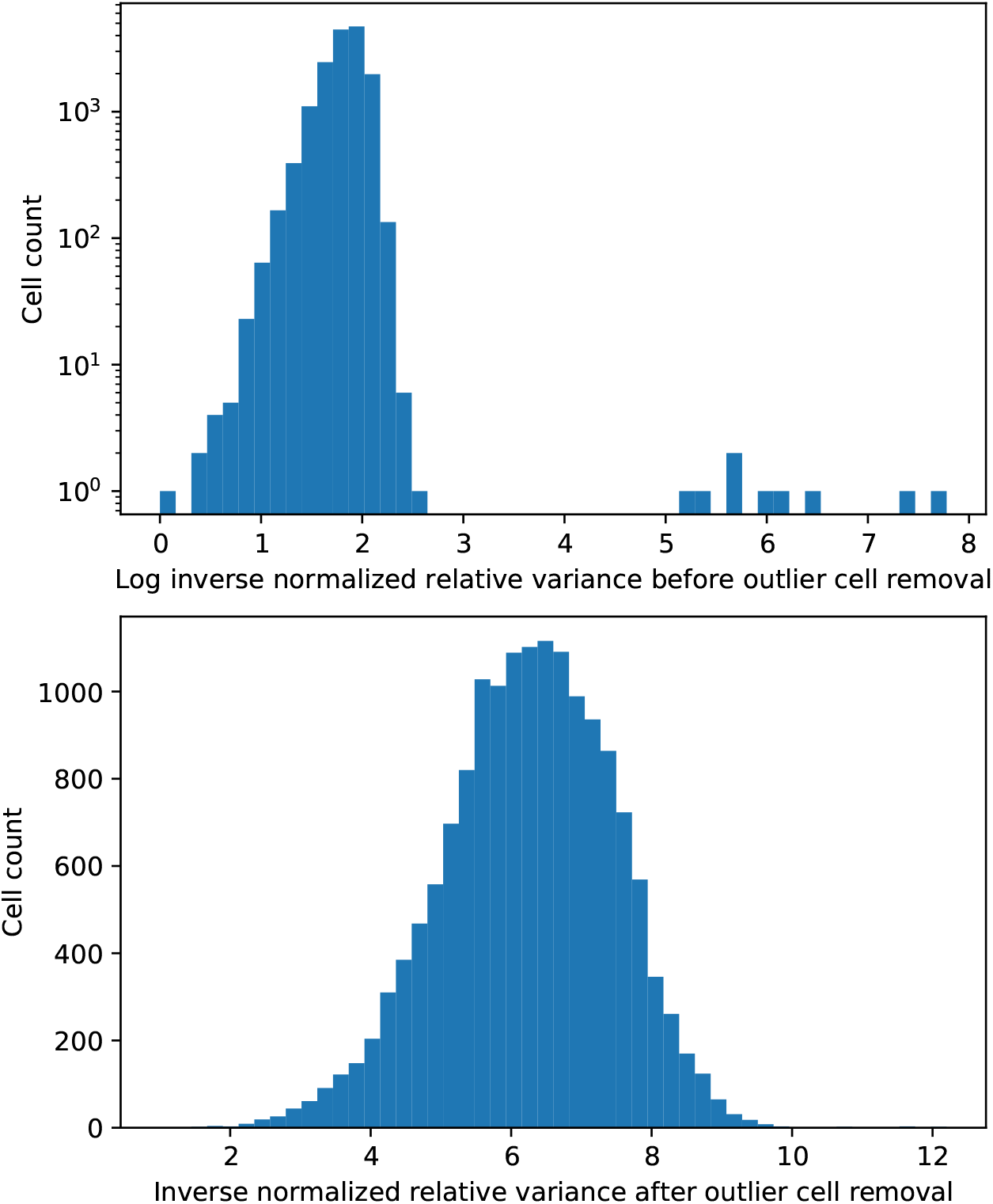
MARS-seq melanoma dataset contained few outlier cells/samples with very low variances. The histograms (Y) show the distributions of inverse normalized relative variances for each cell before in log (X in top) and after in linear (X in bottom) scales for outlier removal in the dysfunctional *v.s*. naive T cell setting. Outlier cells had distinctively lower variances than the major population.

**Supplementary Figure 12:**
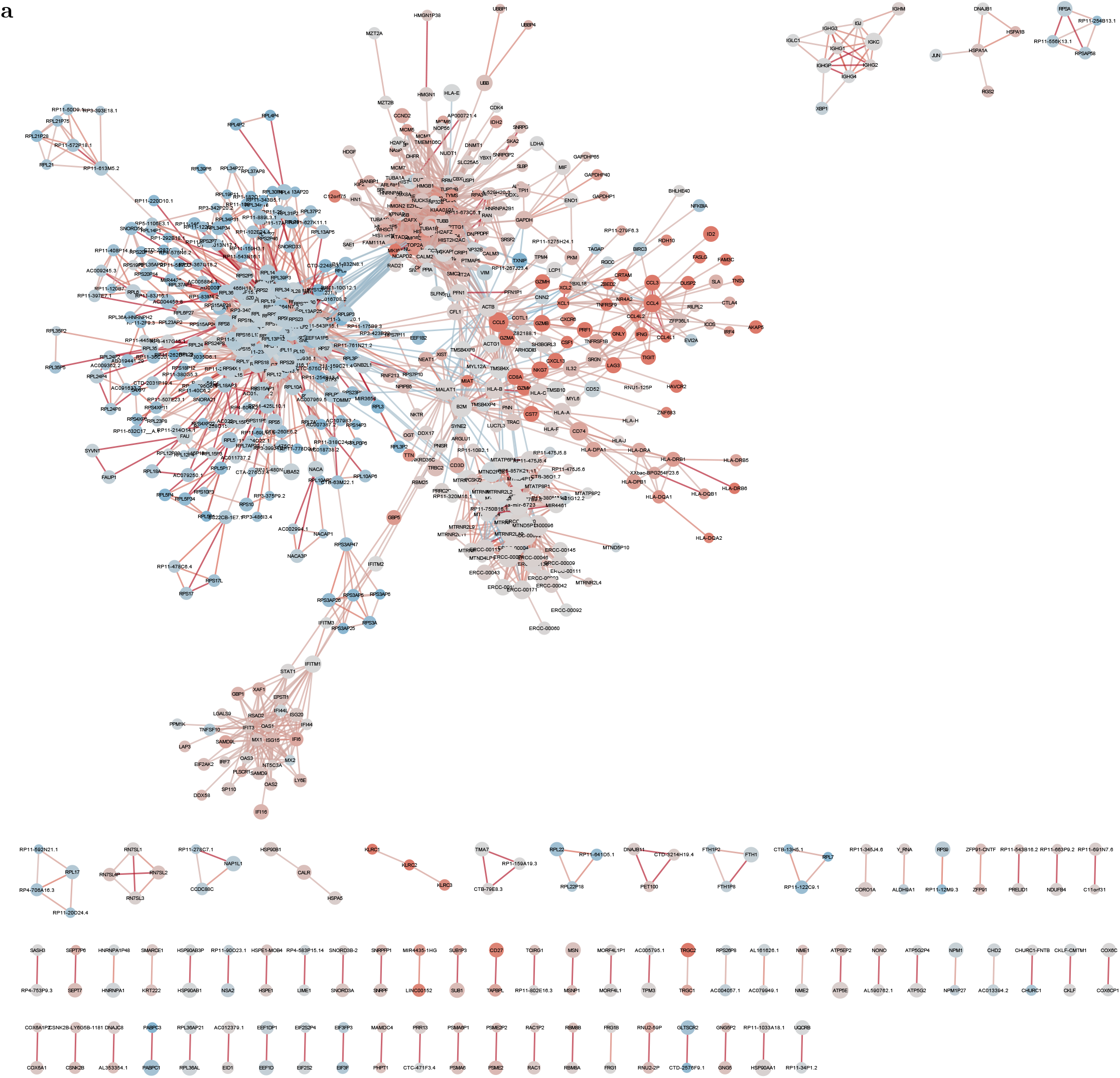
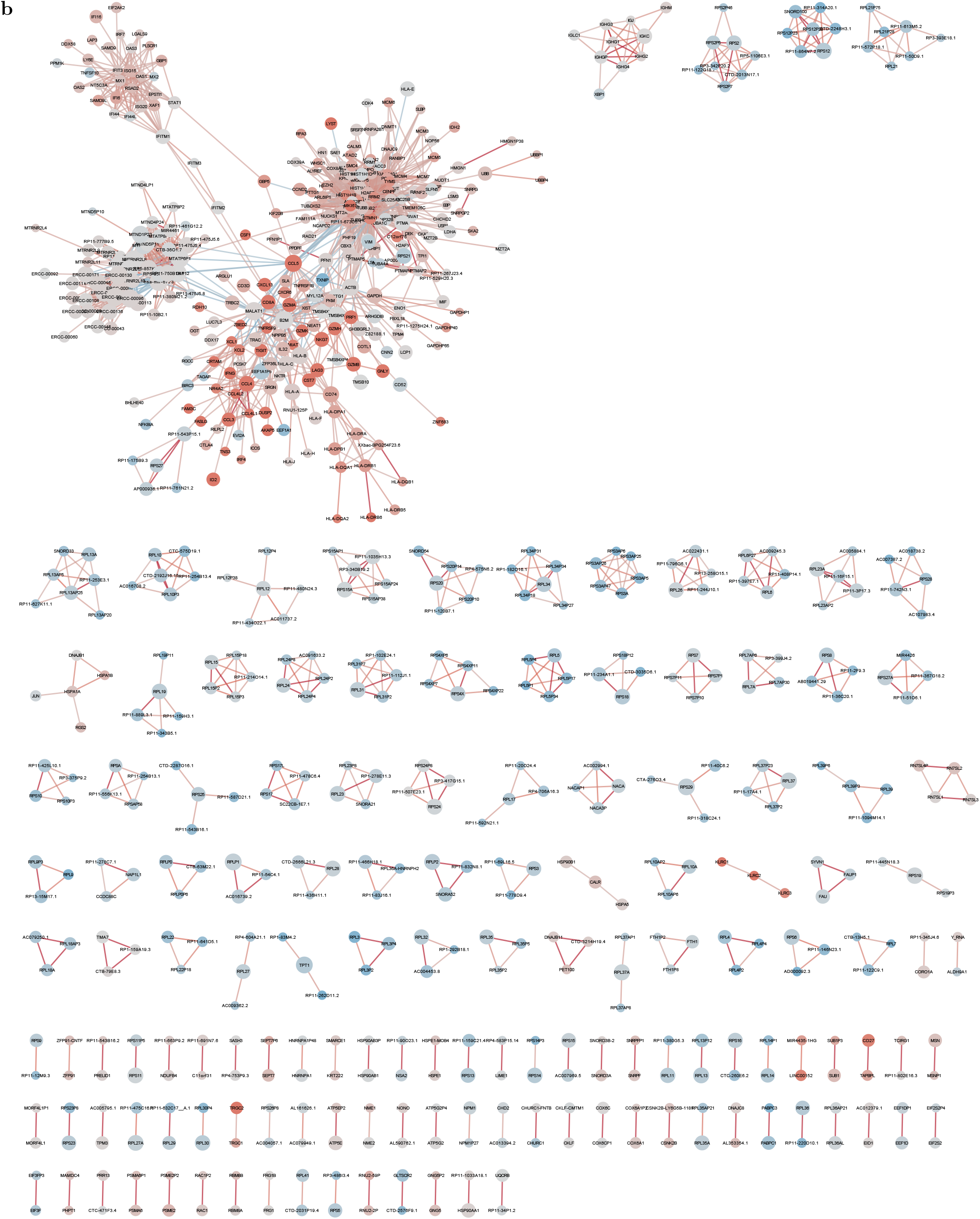
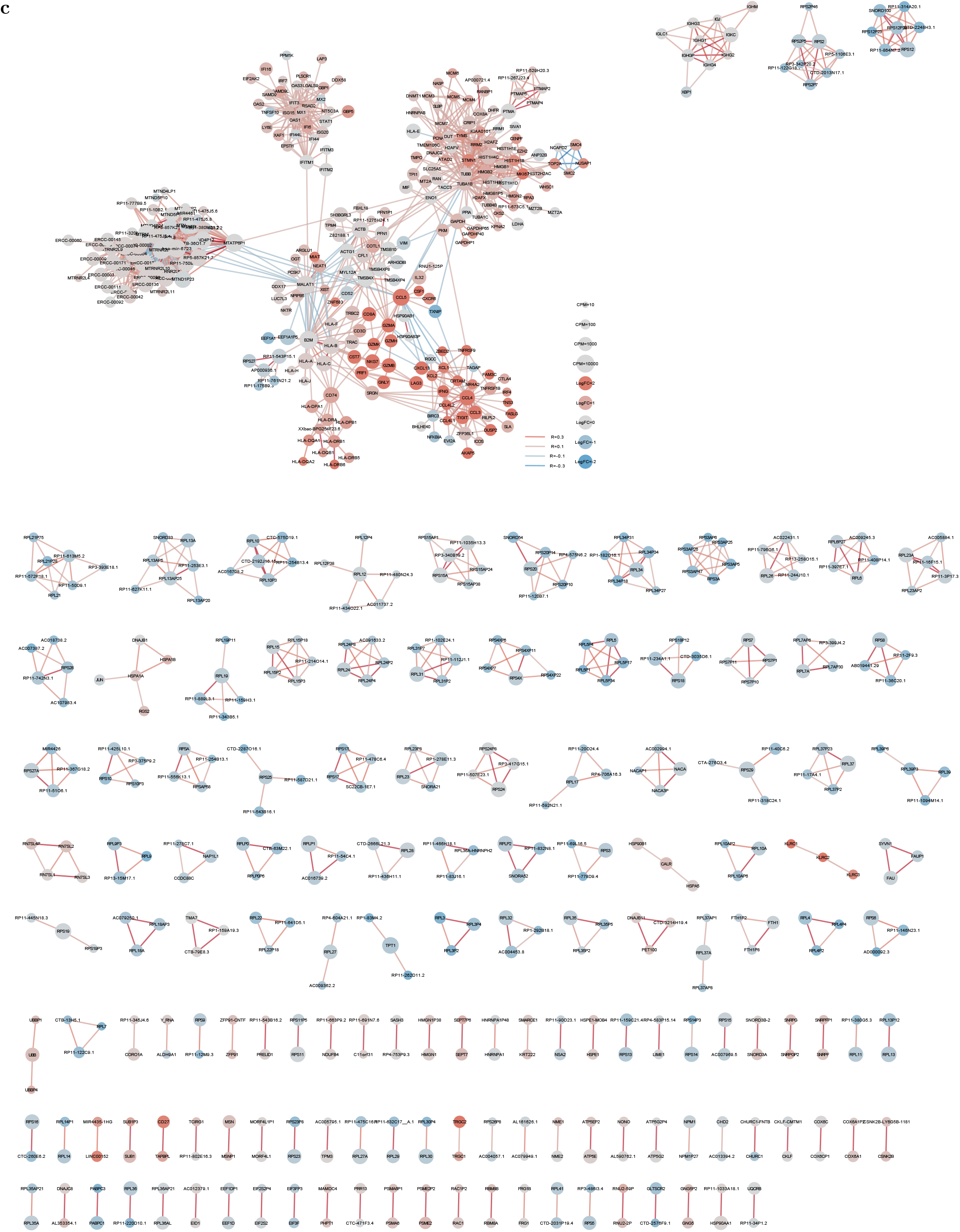
Single-cell transcriptome-wide co-expression networks at each step of GO program removal. Full co-expression networks on the same dataset of dysfunctional T cells from human melanoma at each iterative step of top GO program removal, introducing from 0 to 2 (**a** to **c**) GO covariates. **a** is the original co-expression network. **b** removed cytosolic part (GO:0044445) from **a**. **c** removed chromosome condensation (GO:0030261) from **b**. Fig. 4c further removed the non-coding cluster (left in panel **c**) and the minor connected components from **c**. Legend is in **c**. Zooming in on a digital device is advised.

**Supplementary Figure 13:**
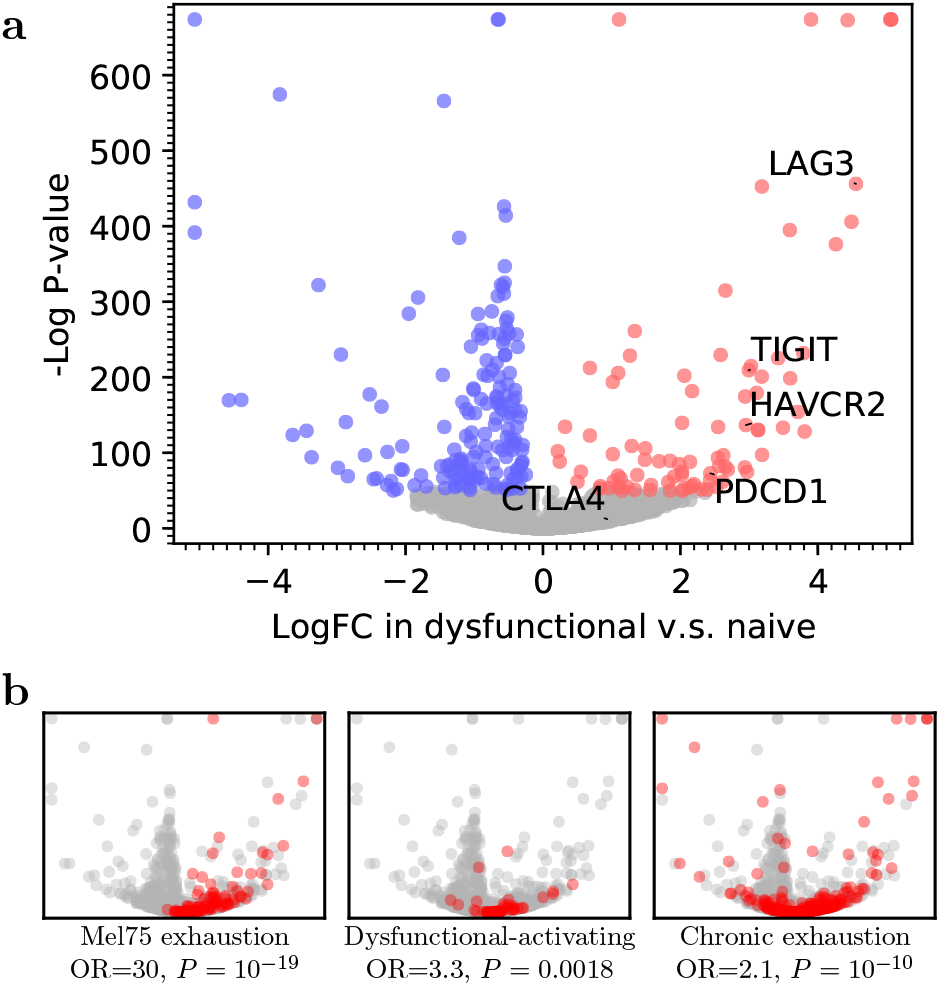
Normalisr identified DEs between dysfunctional and naive T cells in human melanoma MARS-seq. **a** Normalisr detected expression changes associated with T cell dysfunction, shown in a volcano plot of single-cell DE between logFC (X) and −log P-value (Y). Red/blue indicates up-/down-regulated genes (Benjamini-Hochberg *Q*-value ≤ 10^-20^). **b** Up-regulated genes significantly overlapped with published T cell dysfunction signature gene sets (red) in terms of odds ratio (OR) and hypergeometric P-value, including (left-to-right) the Mel75 exhaustion signature in scRNA-seq of human melanoma [38], the dysfunctional-activating signature in bulk RNA-seq of implanted mouse MC38 tumors [36], and the chronic T cell exhaustion signature in bulk RNA-seq of lymphocytic choriomeningitis virus infected mice [37].

Supplementary Table 1: Gene, cell, and gRNA counts of all studies.

Supplementary Table 2: Unexpected associations and potential off-target effects of NTC gRNAs in CROP-seq screen.

Supplementary Table 3: Gene regulatory network inferred from CROP-seq screen.

Supplementary Table 4: Top GO enrichments of inferred targets of top up-/down-regulation favored regulators in CROP-seq screen.

Supplementary Table 5: Significant gene ontology enrichments of top 100 principal genes in the co-expression network of dysfunctional T cells, for different numbers of GO covariates in iterative GO pathway removal.

Supplementary Table 6: Differential gene expression result in dysfunctional *v.s*. naive T cells.

Supplementary Table 7: Node and edge properties of Fig. 4c and Fig. S12.

Supplementary Table 8: Top gene ontology enrichments of the cell cycle and type I interferon response clusters in the co-expression network of dysfunctional T cells.

Supplementary Table 9: Over-abundance of co-expression edges between genes in the same GO or KEGG pathways.

## References

[1] Finak, G., McDavid, A., Yajima, M., Deng, J., Gersuk, V., Shalek, A.K., Slichter, C.K., Miller, H.W., McElrath, M.J., Prlic, M., Linsley, P.S., Gottardo, R.: MAST: a flexible statistical framework for assessing transcriptional changes and characterizing heterogeneity in single-cell RNA sequencing data. Genome Biology 16(1), 278 (2015). doi:10.1186/s13059-015-0844-5. Accessed 2020-02-10

[2] Skinnider, M.A., Squair, J.W., Foster, L.J.: Evaluating measures of association for single-cell transcriptomics. Nature Methods 16(5), 381 (2019). doi:10.1038/s41592-019-0372-4. Accessed 2019-05-06

[3] Mohammadi, S., Davila-Velderrain, J., Kellis, M.: Reconstruction of Cell-type-Specific Interactomes at Single-Cell Resolution. Cell Systems 9(6), 559–5684 (2019). doi:10.1016/j.cels.2019.10.007. Accessed 2020-01-23

[4] Qiu, X., Rahimzamani, A., Wang, L., Ren, B., Mao, Q., Durham, T., McFaline-Figueroa, J.L., Saunders, L., Trapnell, C., Kannan, S.: Inferring Causal Gene Regulatory Networks from Coupled Single-Cell Expression Dynamics Using Scribe. Cell Systems 10(3), 265–27411 (2020). doi:10.1016/j.cels.2020.02.003. Accessed 2020-04-26

[5] Pratapa, A., Jalihal, A.P., Law, J.N., Bharadwaj, A., Murali, T.M.: Benchmarking algorithms for gene regulatory network inference from single-cell transcriptomic data. Nature Methods (2020). doi:10.1038/s41592-019-0690-6. Accessed 2020-02-06

[6] Adamson, B., Norman, T.M., Jost, M., Cho, M.Y., Nuez, J.K., Chen, Y., Villalta, J.E., Gilbert, L.A., Horlbeck, M.A., Hein, M.Y., Pak, R.A., Gray, A.N., Gross, C.A., Dixit, A., Parnas, O., Regev, A., Weissman, J.S.: A Multiplexed Single-Cell CRISPR Screening Platform Enables Systematic Dissection of the Unfolded Protein Response. Cell 167(7), 1867–188221 (2016). doi:10.1016/j.cell.2016.11.048. Accessed 2018-04-23

[7] Gasperini, M., Hill, A.J., McFaline-Figueroa, J.L., Martin, B., Kim, S., Zhang, M.D., Jackson, D., Leith, A., Schreiber, J., Noble, W.S., Trapnell, C., Ahituv, N., Shendure, J.: A Genome-wide Framework for Mapping Gene Regulation via Cellular Genetic Screens. Cell 176(1-2), 377–39019 (2019). doi:10.1016/j.cell.2018.11.029. Accessed 2019-01-14

[8] Soneson, C., Robinson, M.D.: Bias, robustness and scalability in single-cell differential expression analysis. Nature Methods 15(4), 255–261 (2018). doi:10.1038/nmeth.4612. Accessed 2018-07-03

[9] Robinson, M.D., McCarthy, D.J., Smyth, G.K.: edgeR: a Bioconductor package for differential expression analysis of digital gene expression data. Bioinformatics 26(1), 139–140 (2010). doi:10.1093/bioinformatics/btp616. Publisher: Oxford Academic. Accessed 2020-03-02

[10] Townes, F.W., Hicks, S.C., Aryee, M.J., Irizarry, R.A.: Feature selection and dimension reduction for single-cell RNA-Seq based on a multinomial model. Genome Biology 20(1), 295 (2019). doi:10.1186/s13059-019-1861-6. Accessed 2020-05-11

[11] Yang, L., Zhu, Y., Yu, H., Cheng, X., Chen, S., Chu, Y., Huang, H., Zhang, J., Li, W.: scMAGeCK links genotypes with multiple phenotypes in single-cell CRISPR screens. Genome Biology 21(1), 19 (2020). doi:10.1186/s13059-020-1928-4. Accessed 2020-05-11

[12] Katsevich, E., Barry, T., Roeder, K.: Conditional resampling improves calibration and sensitivity in single-cell CRISPR screen analysis. bioRxiv, 2020–0813250092 (2021). doi:10.1101/2020.08.13.250092. Publisher: Cold Spring Harbor Laboratory Section: New Results. Accessed 2021-02-25

[13] Hafemeister, C., Satija, R.: Normalization and variance stabilization of single-cell RNA-seq data using regularized negative binomial regression. Genome Biology 20(1), 296 (2019). doi:10.1186/s13059-019-1874-1. Accessed 2020-02-06

[14] Tang, W., Bertaux, F., Thomas, P., Stefanelli, C., Saint, M., Marguerat, S., Shahrezaei, V.: bayNorm: Bayesian gene expression recovery, imputation and normalization for single-cell RNA-sequencing data. Bioinformatics 36(4), 1174–1181 (2020). doi:10.1093/bioinformatics/btz726. Publisher: Oxford Academic. Accessed 2020-09-04

[15] Breda, J., Zavolan, M., Nimwegen, E.v.: Bayesian inference of the gene expression states of single cells fromscRNA-seqdata. bioRxiv, 2019–1228889956 (2019). doi:10.1101/2019.12.28.889956. Publisher: Cold Spring Harbor Laboratory Section: New Results. Accessed 2020-09-02

[16] Allocco, D.J., Kohane, I.S., Butte, A.J.: Quantifying the relationship between co-expression, co-regulation and gene function. BMC Bioinformatics 5(1), 18 (2004). doi:10.1186/1471-2105-5-18. Accessed 2020-04-29

[17] Loh, P.-R., Tucker, G., Bulik-Sullivan, B.K., Vilhjlmsson, B.J., Finucane, H.K., Salem, R.M., Chasman, D.I., Ridker, P.M., Neale, B.M., Berger, B., Patterson, N., Price, A.L.: Efficient Bayesian mixed-model analysis increases association power in large cohorts. Nature Genetics 47(3), 284–290 (2015). doi:10.1038/ng.3190. Accessed 2018-05-24

[18] Casale, F.P., Rakitsch, B., Lippert, C., Stegle, O.: Efficient set tests for the genetic analysis of correlated traits. Nature Methods 12(8), 755–758 (2015). doi:10.1038/nmeth.3439. Accessed 2017-05-09

[19] Doss, S., Schadt, E.E., Drake, T.A., Lusis, A.J.: Cis-acting expression quantitative trait loci in mice. Genome Research 15(5), 681–691 (2005). doi:10.1101/gr.3216905. Company: Cold Spring Harbor Laboratory Press Distributor: Cold Spring Harbor Laboratory Press Institution: Cold Spring Harbor Laboratory Press Label: Cold Spring Harbor Laboratory Press Publisher: Cold Spring Harbor Lab. Accessed 2020-04-29

[20] Chen, L.S., Emmert-Streib, F., Storey, J.D.: Harnessing naturally randomized transcription to infer regulatory relationships among genes. Genome Biology 8, 219 (2007). doi:10.1186/gb-2007-8-10-r219. Accessed 2017-06-08

[21] Wang, L., Michoel, T.: Efficient and accurate causal inference with hidden confounders from genome-transcriptome variation data. PLOS Computational Biology 13(8), 1005703 (2017). doi:10.1371/journal.pcbi.1005703. Accessed 2017-09-01

[22] Svensson, V.: Droplet scRNA-seq is not zero-inflated. Nature Biotechnology 38(2), 147–150 (2020). doi:10.1038/s41587-019-0379-5. Accessed 2020-04-17

[23] Kim, J.K., Kolodziejczyk, A.A., Ilicic, T., Teichmann, S.A., Marioni, J.C.: Characterizing noise structure in single-cell RNA-seq distinguishes genuine from technical stochastic allelic expression. Nature Communications 6, 8687 (2015). doi:10.1038/ncomms9687. Accessed 2018-05-15

[24] Dijk, D.v., Sharma, R., Nainys, J., Yim, K., Kathail, P., Carr, A.J., Burdziak, C., Moon, K.R., Chaffer, C.L., Pattabiraman, D., Bierie, B., Mazutis, L., Wolf, G., Krishnaswamy, S., Peer, D.: Recovering Gene Interactions from Single-Cell Data Using Data Diffusion. Cell 174(3), 716–72927 (2018). doi:10.1016/j.cell.2018.05.061. Accessed 2018-07-26

[25] Eraslan, G., Simon, L.M., Mircea, M., Mueller, N.S., Theis, F.J.: Single-cell RNA-seq denoising using a deep count autoencoder. Nature Communications 10(1), 390 (2019). doi:10.1038/s41467-018-07931-2. Accessed 2019-07-19

[26] Li, H., van der Leun, A.M., Yofe, I., Lubling, Y., Gelbard-Solodkin, D., van Akkooi, A.C.J., van den Braber, M., Rozeman, E.A., Haanen, J.B.A.G., Blank, C.U., Horlings, H.M., David, E., Baran, Y., Bercovich, A., Lifshitz, A., Schumacher, T.N., Tanay, A., Amit, I.: Dysfunctional CD8 T Cells Form a Proliferative, Dynamically Regulated Compartment within Human Melanoma. Cell 176(4), 775–78918 (2019). doi:10.1016/j.cell.2018.11.043. Accessed 2020-03-17

[27] Jin, X., Simmons, S.K., Guo, A., Shetty, A.S., Ko, M., Nguyen, L., Jokhi, V., Robinson, E., Oyler, P., Curry, N., Deangeli, G., Lodato, S., Levin, J.Z., Regev, A., Zhang, F., Arlotta, P.: In vivo Perturb-Seq reveals neuronal and glial abnormalities associated with autism risk genes. Science 370(6520) (2020). doi:10.1126/science.aaz6063. Publisher: American Association for the Advancement of Science Section: Research Article. Accessed 2020-12-22

[28] Storey, J.D., Tibshirani, R.: Statistical significance for genomewide studies. Proceedings of the National Academy of Sciences 100(16), 9440–9445 (2003). doi:10.1073/pnas.1530509100. Accessed 2017-10-31

[29] Lippert, C., Listgarten, J., Liu, Y., Kadie, C.M., Davidson, R.I., Heckerman, D.: FaST linear mixed models for genome-wide association studies. Nature Methods 8(10), 833–835 (2011). doi:10.1038/nmeth.1681. Accessed 2018-05-24

[30] Meola, N., Domanski, M., Karadoulama, E., Chen, Y., Gentil, C., Pultz, D., Vitting-Seerup, K., Lykke-Andersen, S., Andersen, J., Sandelin, A., Jensen, T.: Identification of a Nuclear Exosome Decay Pathway for Processed Transcripts. Molecular Cell 64(3), 520–533 (2016). doi:10.1016/j.molcel.2016.09.025. Accessed 2021-03-14

[31] Strating, J.R.P.M., Martens, G.J.M.: The p24 family and selective transport processes at the ERGolgi interface. Biology of the Cell 101(9), 495–509 (2009). doi:10.1042/BC20080233. _eprint: https://onlinelibrary.wiley.com/doi/pdf/10.1042/BC20080233. Accessed 2020-03-02

[32] Morgens, D.W., Deans, R.M., Li, A., Bassik, M.C.: Systematic comparison of CRISPR/Cas9 and RNAi screens for essential genes. Nature Biotechnology 34(6), 634–636 (2016). doi:10.1038/nbt.3567. Accessed 2019-07-22

[33] Stenmark, H., Olkkonen, V.M.: The Rab GTPase family. Genome Biology 2(5), 3007–1 (2001). doi:10.1186/gb-2001-2-5-reviews3007. Accessed 2020-03-02

[34] An integrated encyclopedia of DNA elements in the human genome. Nature 489(7414), 57–74 (2012). doi:10.1038/nature11247. Number: 7414 Publisher: Nature Publishing Group. Accessed 2020-04-09

[35] Davis, C.A., Hitz, B.C., Sloan, C.A., Chan, E.T., Davidson, J.M., Gabdank, I., Hilton, J.A., Jain, K., Baymuradov, U.K., Narayanan, A.K., Onate, K.C., Graham, K., Miyasato, S.R., Dreszer, T.R., Strattan, J.S., Jolanki, O., Tanaka, F.Y., Cherry, J.M.: The Encyclopedia of DNA elements (ENCODE): data portal update. Nucleic Acids Research 46(D1), 794–801 (2018). doi:10.1093/nar/gkx1081. Publisher: Oxford Academic. Accessed 2020-04-09

[36] Singer, M., Wang, C., Cong, L., Marjanovic, N.D., Kowalczyk, M.S., Zhang, H., Nyman, J., Sakuishi, K., Kurtulus, S., Gennert, D., Xia, J., Kwon, J.Y.H., Nevin, J., Herbst, R.H., Yanai, I., Rozenblatt-Rosen, O., Kuchroo, V.K., Regev, A., Anderson, A.C.: A Distinct Gene Module for Dysfunction Uncoupled from Activation in Tumor-Infiltrating T Cells. Cell 166(6), 1500–15119 (2016). doi:10.1016/j.cell.2016.08.052. Publisher: Elsevier. Accessed 2020-03-19

[37] Doering, T., Crawford, A., Angelosanto, J., Paley, M., Ziegler, C., Wherry, E..: Network Analysis Reveals Centrally Connected Genes and Pathways Involved in CD8+ T Cell Exhaustion versus Memory. Immunity 37(6), 1130–1144 (2012). doi:10.1016/j.immuni.2012.08.021. Accessed 2020-03-25

[38] Tirosh, I., Izar, B., Prakadan, S.M., Wadsworth, M.H., Treacy, D., Trombetta, J.J., Rotem, A., Rodman, C., Lian, C., Murphy, G., Fallahi-Sichani, M., Dutton-Regester, K., Lin, J.-R., Cohen, O., Shah, P., Lu, D., Genshaft, A.S., Hughes, T.K., Ziegler, C.G.K., Kazer, S.W., Gaillard, A., Kolb, K.E., Villani, A.-C., Johannessen, C.M., Andreev, A.Y., Allen, E.M.V., Bertagnolli, M., Sorger, P.K., Sullivan, R.J., Flaherty, K.T., Frederick, D.T., Jan-Valbuena, J., Yoon, C.H., Rozenblatt-Rosen, O., Shalek, A.K., Regev, A., Garraway, L.A.: Dissecting the multicellular ecosystem of metastatic melanoma by single-cell RNA-seq. Science 352(6282), 189–196 (2016). doi:10.1126/science.aad0501. Publisher: American Association for the Advancement of Science Section: Research Article. Accessed 2020-03-25

[39] Rebhan, M., Chalifa-Caspi, V., Prilusky, J., Lancet, D.: GeneCards: a novel functional genomics compendium with automated data mining and query reformulation support. Bioinformatics 14(8), 656–664 (1998). doi:10.1093/bioinformatics/14.8.656. Accessed 2021-03-26

[40] Thommen, D.S., Schumacher, T.N.: T Cell Dysfunction in Cancer. Cancer Cell 33(4), 547–562 (2018). doi:10.1016/j.ccell.2018.03.012. Accessed 2020-05-18

[41] Crow, M., Paul, A., Ballouz, S., Huang, Z.J., Gillis, J.: Exploiting single-cell expression to characterize co-expression replicability. Genome Biology 17(1), 101 (2016). doi:10.1186/s13059-016-0964-6. Accessed 2020-09-02

[42] Lhnemann, D., Kster, J., Szczurek, E., McCarthy, D.J., Hicks, S.C., Robinson, M.D., Vallejos, C.A., Campbell, K.R., Beerenwinkel, N., Mahfouz, A., Pinello, L., Skums, P., Stamatakis, A., Attolini, C.S.-O., Aparicio, S., Baaijens, J., Balvert, M., Barbanson, B.d., Cappuccio, A., Corleone, G., Dutilh, B.E., Florescu, M., Guryev, V., Holmer, R., Jahn, K., Lobo, T.J., Keizer, E.M., Khatri, I., Kielbasa, S.M., Korbel, J.O., Kozlov, A.M., Kuo, T.-H., Lelieveldt, B.P.F., Mandoiu, I.I., Marioni, J.C., Marschall, T., Mlder, F., Niknejad, A., Raczkowski, L., Reinders, M., Ridder, J.d., Saliba, A.-E., Somarakis, A., Stegle, O., Theis, F.J., Yang, H., Zelikovsky, A., McHardy, A.C., Raphael, B.J., Shah, S.P., Schnhuth, A.: Eleven grand challenges in single-cell data science. Genome Biology 21(1), 31 (2020). doi:10.1186/s13059-020-1926-6. Accessed 2020-04-28

[43] Basso, K., Margolin, A.A., Stolovitzky, G., Klein, U., Dalla-Favera, R., Califano, A.: Reverse engineering of regulatory networks in human B cells. Nature Genetics 37(4), 382–390 (2005). doi:10.1038/ng1532. Accessed 2017-05-09

[44] Scutari, M.: Learning Bayesian Networks with the bnlearn R Package. Journal of Statistical Software 35(3) (2010). doi:10.18637/jss.v035.i03. Accessed 2018-02-22

[45] Barkas, N., Petukhov, V., Nikolaeva, D., Lozinsky, Y., Demharter, S., Khodosevich, K., Kharchenko, P.V.: Joint analysis of heterogeneous single-cell RNA-seq dataset collections. Nature Methods 16(8), 695–698 (2019). doi:10.1038/s41592-019-0466-z. Accessed 2019-08-02

[46] Li, W.V., Li, J.J.: An accurate and robust imputation method scImpute for single-cell RNA-seq data. Nature Communications 9(1), 997 (2018). doi:10.1038/s41467-018-03405-7. Accessed 2018-06-15

[47] Risso, D., Perraudeau, F., Gribkova, S., Dudoit, S., Vert, J.-P.: A general and flexible method for signal extraction from single-cell RNA-seqdata. Nature Communications 9(1), 1–17 (2018). doi:10.1038/s41467-017-02554-5. Number: 1 Publisher: Nature Publishing Group. Accessed 2020-03-02

[48] Lopez, R., Regier, J., Cole, M.B., Jordan, M.I., Yosef, N.: Deep generative modeling for single-cell transcriptomics. Nature Methods 15(12), 1053 (2018). doi:10.1038/s41592-018-0229-2. Accessed 2018-12-07

[49] Vieth, B., Ziegenhain, C., Parekh, S., Enard, W., Hellmann, I.: powsimR: power analysis for bulk and single cell RNA-seq experiments. Bioinformatics 33(21), 3486–3488 (2017). doi:10.1093/bioinformatics/btx435. Publisher: Oxford Academic. Accessed 2020-04-30

[50] Klopfenstein, D.V., Zhang, L., Pedersen, B.S., Ramrez, F., Vesztrocy, A.W., Naldi, A., Mungall, C.J., Yunes, J.M., Botvinnik, O., Weigel, M., Dampier, W., Dessimoz, C., Flick, P., Tang, H.: GOATOOLS: A Python library for Gene Ontology analyses. Scientific Reports 8(1), 1–17 (2018). doi:10.1038/s41598-018-28948-z. Number: 1 Publisher: Nature Publishing Group. Accessed 2020-03-18

[51] Strimmer, K.: fdrtool: a versatile R package for estimating local and tail area-based false discovery rates. Bioinformatics 24(12), 1461–1462 (2008). doi:10.1093/bioinformatics/btn209. Accessed 2017-10-31

[52] Cokelaer, T., Pultz, D., Harder, L.M., Serra-Musach, J., Saez-Rodriguez, J.: BioServices: a common Python package to access biological Web Services programmatically. Bioinformatics 29(24), 3241–3242 (2013). doi:10.1093/bioinformatics/btt547. Publisher: Oxford Academic. Accessed 2020-04-13

